# A new thermophysiological model unifying heat balance theory and physiological feedback control: *Framework, heat defence functions, and individualisation*

**DOI:** 10.64898/2026.11.20.768055

**Authors:** Josh Foster, Richard S. Whittle, Shoji Leach, Ellie J. Allsop, Andrew D’Souza, Peter D. Hodkinson, Sean R. Notley, Zachary J. McKenna, Hannah Pallubinsky, Simon G. Hodder, Craig G. Crandall, Glen P. Kenny, Andrej A. Romanovsky, Robert D Meade

## Abstract

Coded in Julia using ModelingToolkit, we developed a new computational model (TeMP) that calculates the emergent physiological responses to heat with and without physical activity (PA). Rooted in established physical and physiological principles, we detail the model framework, major novelties, and sources of individual variation. TeMP uses physical heat-transfer and heat-balance concepts to determine the rate of whole-body heat storage (S) over a given exposure period, based on the dynamic relationship between S, tissue temperature, and thermoeffector function. Each thermoeffector (skin blood flow, sweating, and ventilation) operates on independent feedback loops and is modelled based on afferent inputs from central and peripheral/skin tissue temperatures. All thermoeffectors use simple piecewise equations to determine the onset threshold, slope, and maximum functions, removing the need for a central error signal pervasive in alternative models. Skin blood flow is bounded by individual fitness level and is attenuated by PA, whereas sweating is more sensitive during PA due to heightened sympathetic activity. Minute ventilation is also bounded by individual fitness and is sensitive to body temperature and PA intensity, capturing the disproportionate increase in breathing rate at the ventilatory threshold. Thermoeffector functions are individualised for intrinsic characteristics, including heat acclimatisation, age, menstrual cycle phase, skin grafting, sympathetic blockade (i.e., spinal cord lesion), and severe energy restriction (i.e., anorexia nervosa). Finally, heart and rectal temperature are modelled independently and follow similar divergent patterns shown in empirical studies at rest and during PA.

## Background

Accurate prediction of physiological outcomes during heat stress can inform policy decisions and targeted interventions. Although lab- or field-based empirical studies are considered the gold standard method to generate such insights, they are expensive, time-consuming, and are generally considered high-risk procedures, especially when conducted in heat-vulnerable groups. Computational models, on the other hand, can provide useful insights into the likely magnitude and time course of physiological strain and risk stratification associated with extreme heat exposure. Physiological models, therefore, can function as a decision support tool for identifying at risk groups and contexts, triaging potential adaptive strategies, and designing safe but effective laboratory or field-based studies. While models do exist for these purposes, they carry fundamental issues that limit their applicability. Three primary issues include: an outdated physiological control architecture, poor scope for individualisation, and limited ability to model heat-defence responses during physical activity.

Coded in Julia using *ModelingToolkit*, we developed TeMP, a dynamic physiological model of human thermoregulation. Unlike previous models built from an outdated error-control framework (Stolwijk & Hardy, 1966), our model requires no arbitrary reference temperature (“set point”), and effector functions are not built to fit a single dataset in isolation. TeMP instead explicitly integrates current understanding of biophysical and physiological principles, which dictate that steady-state body temperature (*T*_body_) emerges as the equilibrium point (or balance point) where whole-body heat gain is matched by an appropriately weighted physiological response required to arrest the rate of whole-body heat storage (Werner, 2010; Meade *et al*., 2024b). Independent thermoeffector functions controlling both heat and cold defences not only better reflect current understanding of the human thermoregulatory architecture (Romanovsky, 2007; Werner, 2010; Romanovsky, 2018), but also means that almost all components can be modified to better reflect variation in thermoregulatory function due to any individual factor.

This paper serves as an introduction to TeMP, clarifying its most novel features. The validation and parameter tuning is underway and will be covered in a follow-up manuscript, allowing us to first introduce the core principles TeMP is based on (heat balance theory and physiological feedback control), and merge them into an intuitive physiological model. Against the backdrop of our rapidly warming global climate, this manuscript will focus exclusively on the heat-defence functions (with or without physical activity, PA).

### Overview of heat storage and physiological feedback control

Broadly speaking, thermoregulation can be divided into physical and physiological domains. The physical arm details exchange of thermal energy between body tissues and between the body and the environment. The physiological arm actively modulates these exchanges through physiological (primarily autonomic) thermoeffector responses to achieve heat balance and arrest the development of a critically high *T*_body_ (Romanovsky, 2018). The physiological thermoeffector responses to heat include cutaneous vasodilation, region specific non-cutaneous vasoconstriction, eccrine sweating, and hyperventilation. A key limitation of extant thermophysiological models is that they are often built around only one domain, either physical or physiological, with minimal attention paid to the other, and to how the two domains connect. The aim of this section is to define the theoretical underpinnings and current understanding of how biophysics and physiology integrate, explaining how a given *T*_body_ emerges. Antecedent to this discussion, it is important to state that current evidence does not support the commonly held misconception that there need be a unified control system with a single central controller and a temperature set point that this the thermoregulatory system actively attempts to defend.

### Biophysics of human heat exchange and storage

The goal of the thermoregulatory system is to maintain a state of thermal balance, such that heat gain (from metabolism and potentially the environment) and heat loss are matched, resulting in an equilibrium *T*_body_ optimal for human biochemical function. The heat balance equation can be shown as:

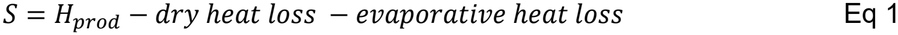

With all units in Watts, *S* is whole body heat storage, *H*_prod_ is metabolic heat production. The Pontzer equation models resting *H*_prod_ from humans aged 8 days to 95 years with fat-free mass as an independent modifier (Pontzer *et al*., 2021). Dry heat loss is attributed to heat transfer that relies chiefly on temperature gradients between the skin and the environment and occurs via radiation, convection, and conduction. Dry heat transfer can push heat toward the body, making the value negative and raising *S*. Evaporative describes heat transfer through pressure gradients, i.e., fluid evaporation from the skin and the respiratory tract epithelia to the environment.

During quiet rest in a cool environment, heat loss to the environment approximates *H*_prod_, resulting in an *S* approaching 0 W (heat balance). Thus, even in ‘thermoneutral’ environments, heat defence is an active physiological process required to prevent excessive thermal energy accumulation. However, the extent to which thermoeffectors are activated can rise sharply with heat stress or PA.

#### Transferring metabolic heat to the skin

*H*_prod_ from cellular metabolism in all organs and tissues in the body, including the viscera in the deep core, is transported to the skin by convection and conduction. Convective heat transport through the blood supply is distributed in a closed loop between the heart, core, muscles, fat, and skin. During heat stress, skin blood flow can heavily increase through cutaneous vasodilation, whereas core flow typically decreases through core (mostly gastrointestinal) vasoconstriction (Minson *et al*., 1998). Conduction between body compartments is a slow form of heat transfer inadequate for human thermal stability in isolation but plays an important role in passive heat transfer from the core to the skin. Figure 1A demonstrates how heat is transferred between each compartment by blood flow and between compartments.

**Figure 1.**
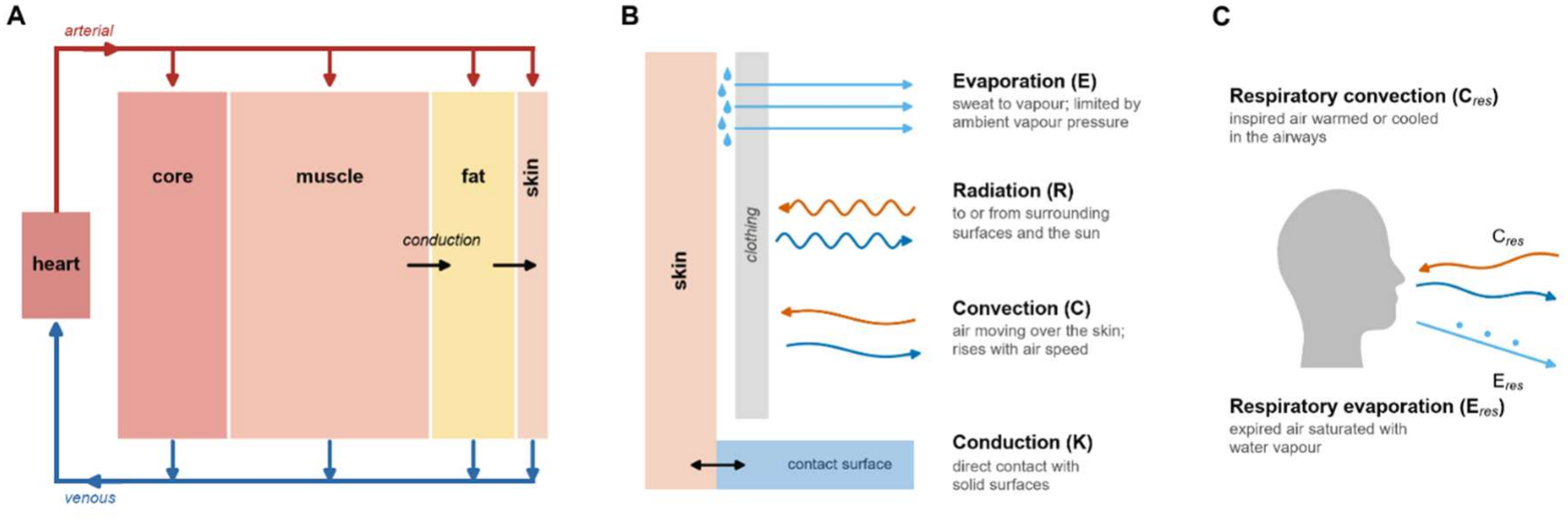
Schematic representation of the primary physical aspects of heat transfer. Panel A shows the pathways by which heat is transferred between body compartments. ‘Core’ is a term used to collapse the deep tissues such as the viscera, bone, and other organs. Core (particularly bone) to skeletal muscle conductive heat transfer is omitted due non-uniformity, which presents various modelling challenges. Convective heat transfer is driven by arterial blood flow delivery to each tissue, heat transfer between the arterial blood and tissue, and venous return to the heart. Panel B shows the pathways by which heat is exchanged between the skin surface and the environment. Bi-directional arrows suggest that heat exchange can occur in either direction, which is true for all ‘dry’ heat transfer terms. Panel C shows the convective and evaporative pathways of respiratory heat exchange.

#### Skin surface heat transfer

Heat arriving at the skin surface from deeper tissues via conduction and convection is further transferred to the environment. Heat transfer between the skin and the environment is governed by fundamental physical laws, which renders modelling straight-forward. The direction of dry heat transfer (depicted in Figure 1B) depends on whether the skin is hotter than the ambient and mean radiant temperature. Conduction depends on the conductivity and temperature of the material, and the surface area of exposed skin to that material.

Respiratory convection also contributes to dry heat transfer but is instead based on the temperature difference between the air and the epithelial lining of the respiratory tract. Respiratory evaporative heat loss is driven by the water vapor pressure gradient, but since the lungs are assumed fully saturated with water vapor, it depends solely on the humidity of the ambient environment. Respiratory heat loss is typically small at rest but becomes relevant at higher activity levels with substantially increased ventilation. Clothing insulation and air velocity impact nearly every avenue of heat transfer at the skin surface and must be considered in thermoregulation models.

Similar to how evaporative heat loss from the respiratory epithelia is driven the surface-environment water vapor pressure gradient, evaporative heat loss at the skin is also determined primarily by the water vapor pressure gradient but, in this case, between the skin surface and the environment. An additional factor affecting evaporative heat loss at the skin surface is the sweat efficiency, which can be determined as the ratio between sweat evaporation (liberate 2.4 kJ of heat per 1 g of sweat) and dripping (liberates no heat).

Efficiency is determined by the sweat rate and the maximum evaporative capacity of the environment. The former implies a physiological limit, while the latter is impacted by clothing, humidity, and air velocity. It is also relevant to consider hidromeiosis, the phenomenon of a reduced sweat output with prolonged wet skin due to blockage of the underlying sweat glands (Candas *et al*., 1983). This effect is especially relevant in hot humid environments and appears to limit wasteful fluid loss. Figure 1B and 1C show the avenues where heat exchange can occur at the skin surface and respiratory tract, respectively.

#### Dynamic whole body heat storage

Whole body heat storage (*S*) changes dynamically throughout any exposure based on interactions between *H*_prod_, dry, and evaporative heat exchange. Immediately upon transition into heat and/or initiation of PA, *S* is positive, with heat being stored at a greater rate than it is lost to the environment. Considering passive physics only (*H*_prod_ + dry), *T*_body_ will equilibrate at a point where dry heat loss matches *H*_prod_, meaning the equilibrium *T*_body_ would typically exceed 40 °C in warm climates. However, adaptive physiological reactions arrest the development of a critically high *T*_body_, because as *T*_body_ rises above a threshold, active skin vasodilation and sweating augment both dry and evaporative heat losses, respectively. Hence, thermophysiological models typically include both a passive physics and active physiological component.

In the next section ‘***Physiological feedback control***’, we briefly discuss the architecture of the human thermoregulatory system, drawing on concepts previously described (Werner, 2010; Romanovsky, 2018). We then integrate the physical and physiological arms to demonstrate how a given *T*_body_ emerges over time.

### Physiological feedback control

The mammalian thermoregulatory system represents a federation of independent thermoeffector loops, each being driven by a unique combination of deep (core) and superficial (shell) temperatures (Romanovsky, 2018). These signals originate from peripheral and central thermoreceptors. Peripheral receptors are located on free endings of sensory neurons predominantly in the skin and mucosa (Jänig, 2018; Romanovsky, 2018), and at-least some of these receptors are likely to be represented by transient receptor potential (TRP) cation channels. Figure 2 illustrates some of the body’s molecular arsenal for detecting physiologically relevant temperatures and thus serving as a signal for various thermoeffectors (Romanovsky, 2018). The activation of warm- and hot-sensitive TRP channels is dependent on the prevailing temperature the channel is exposed to, whereas cold channel activity may also depend in the rate of temperature change (Ran *et al*., 2016). Central thermosensitivity involves warm-sensitive neurons in the preoptic anterior hypothalamus (POA). Mechanisms of central thermosensitivity are likely to include temperature-dependent potassium conductances (K+ currents), synaptic modulation, and nonselective cation currents (Conti, 2018), with TRPM2-mediated increased synaptic drive to the POA warm-sensitive neurons being an example (Kamm *et al*., 2021).

**Figure 2.**
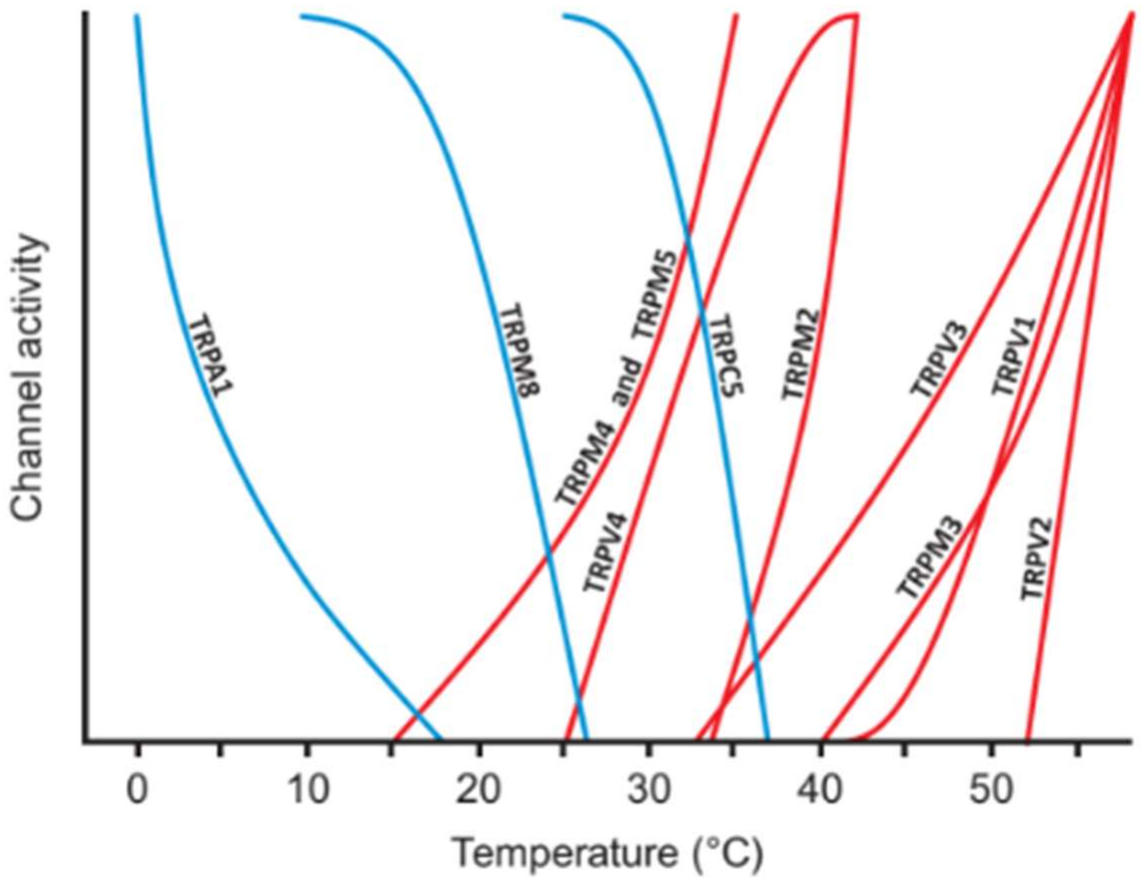
Temperature dependence of thermal TRP channel activity. Note that warm sensitive channels are active in the typical resting deep body temperature range (36.0-37.5°C), meaning that heat loss is constitutively ‘active’ without overt physical activity or environmental heating. Heat loss must be active at rest to counteract the heat produced from resting metabolism, resulting in a stable body temperature and zero net heat storage (*S*). Used with permission from Romanovsky (2018).

For all autonomic thermoeffectors, neural pathways go through the POA (Tabarean, 2018). Neural pathways descending from these neurons provide a POA temperature signal that contributes to driving thermoeffectors such as sweating and skin blood flow (McAllen & McKinley, 2018). The earlier unified-control concepts of thermoregulation (Hammel *et al*., 1963) mimicked engineering models and required neural circuitry comprising 1) an integrator for calculating a *T*_body_, 2) a comparator for comparing this mean *T*_body_ with the set-point *T*_body_, and 3) a coordinator of different thermoeffector responses based on the difference between set-point and actual *T*_body_. These specialised neural circuits were thought to reside in the POA, but none were identified. When intrinsically simple and biologically plausible thermoregulation concepts emerged, the need for a set-point comparator in these specialised circuits disappeared (Romanovsky, 2007; Werner, 2010; Romanovsky, 2018).

In addition, the POA receives inputs from outside of the thermoregulation system which modulate the activity of thermoregulatory circuits (Saper *et al*., 2012). Such non-thermal modulation becomes relevant when considering multiple conditions such as ageing, menstrual phase, energy restriction, and heat acclimatisation. For example, ageing and progesterone (associated with the mid-luteal phase of menstrual cycle) reduce heat-defence thermoeffector activity for any given *T*_body_ (Horvath & Drinkwater, 1982; Charkoudian & Stachenfeld, 2016), while heat acclimatisation increases heat loss for any given *T*_body_ (Havenith, 2001).

#### Unifying heat balance and physiological feedback to generate an emergent body temperature

The previous sections detail the physical and physiological aspects of heat transfer. To determine how *T*_body_ emerges at the intersection of biophysics and physiological feedback control, the workflow reduces to 1) PA and/or environmental heat raises *S* through increased *H*_prod_ or diminished/reversed dry heat loss, respectively, 2) raised *S* increases thermal energy accumulation, proportionally raising *T*_body_, 3) the rise in *T*_body_ is sensed by thermoreceptors which evoke physiological heat loss responses proportional to the absolute *T*_body_ and non-thermoregulatory neural inputs, 4) thermoeffector activity decreases the *S* resulting in either 5) a stabilisation of *T*_body_ occurs once thermoeffector response are sufficient to return *S* to zero (compensable heat stress) or 6) a continuous increase in *T*_body_ if thermoeffector responses are insufficient or if the environment cannot accommodate sufficient dry or evaporative heat loss (uncompensable heat stress).

We reiterate that *T*_body_ is the *signal* that drives the physiological feedback loops required for heat balance, consistent with empirical observations in humans (Meade *et al*., 2023; Meade *et al*., 2024a). This framework diverges from alternative thermoregulation models such as Fiala and JOS-3 (Stolwijk & Hardy, 1966; Fiala *et al*., 2001; Kobayashi & Tanabe, 2013) because they treat the *deviation* from a set-point *T*_body_ as the regulated signal, an outdated concept (Romanovsky, 2018). Interestingly, it has been established that absolute *T*_body_, rather than Δ *T*_body_, represents the regulated variable in a variety of animal models of fever; in contrast, Δ*T*_body_ varies widely in all these fevers and cannot be used as a reliable measure of fever height (Ivanov & Romanovsky, 2002). Figure 3 shows pathways leading to compensable and uncompensable heat stress (Figure 3A), and how each scenario would impact the emergent temperature (Figure 3B). The pathway shown in Figure 3 and described above mirrors TeMP’s architecture and underlying philosophy, abandoning the concept of a thermoregulatory setpoint in favor of a balance point, (Werner, 2010; Romanovsky, 2018), which more accurately resembles current understanding of how the human thermoregulatory system functions..

**Figure 3.**
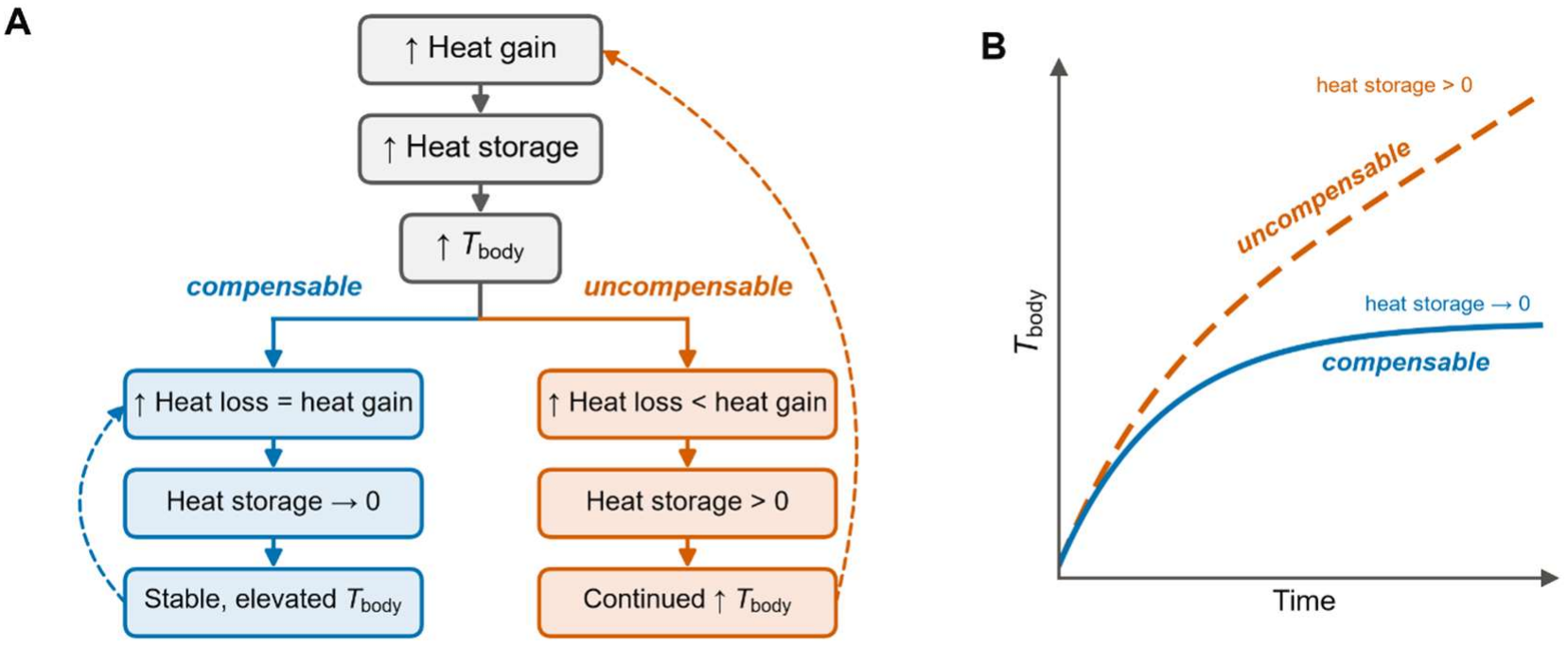
Simplified framework leading to an emergent body temperature without the need for a set-point temperature. Panel A shows a compensable and uncompensable exposure pathway. In the compensable pathway, heat loss is sufficient to offset the heat gain, bring heat storage to zero, and stabilise body temperature. In the uncompensable pathway, heat loss is insufficient to bring heat storage to zero, resulting in a continuous increase in body temperature. Sources of inadequate heat loss can be driven by humidity, clothing, or inadequate sweating caused by intrinsic factors (i.e., ageing, medication, dehydration, skin injuries etc). Panel B shows how each exposure pathway translates into the emergent body temperature throughout the series of a given exposure.

### Novel aspects of TeMP

TeMP adopts the architecture described in the previous section. The chosen architecture allows TeMP’s effector control to be built from physiological principles rather than regression fitted to whole-body outcomes. Following appropriate calibration and independent validation, TeMP may provide a platform for examining how individual characteristics influence predicted thermal responses across a range of environmental and physical activity conditions. Building on our discussion of the fundamental biophysical and physiological concepts TeMP is based on, here we describe the most pertinent aspects of TeMP that distinguishes it from existing models.

### Decentralised framework

Our framework integrates heat balance concepts and physiological feedback control (Figure 4) without the need for a set-point controller. First, heat is ‘delivered’ into various body compartments (nodes) through metabolism (*H*_prod_), where it disperses through convection and conduction between seven distinct compartments including the heart, lumped core, rectum, active and inactive skeletal muscle, fat, and skin. The passive system’s modifiable components include resting *H*_prod_, body type, and clothing insulation Thermal and evaporative resistance of clothing is modelled based on ISO 9920. During PA, additional *H*_prod_ is delivered to the active skeletal muscle, where its proportion to inactive muscle depends on the PA type (walking, running, or cycling with or without load carriage). For walking and running, the additional *H*_prod_ with load carriage is derived from the ACSM metabolic equations (ACSM, 1986), in which the generic resting term (3.5 mL·kg^−1^·min^−1^) is replaced by the individual’s Pontzer-derived resting *H*_prod_ (Pontzer *et al*., 2021), so that only the activity component of the equation is retained. Carried load (*L*) is included by applying the equations to total carried mass, scaling 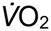 by 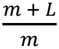 (Goldman & Iampietro, 1962), which agrees with the Pandolf load carriage equation (Pandolf *et al*., 1977) to within ~5% for loads up to 40 kg. For cycling, *H*_prod_ is derived from external power assuming 20% gross efficiency.

**Figure 4.**
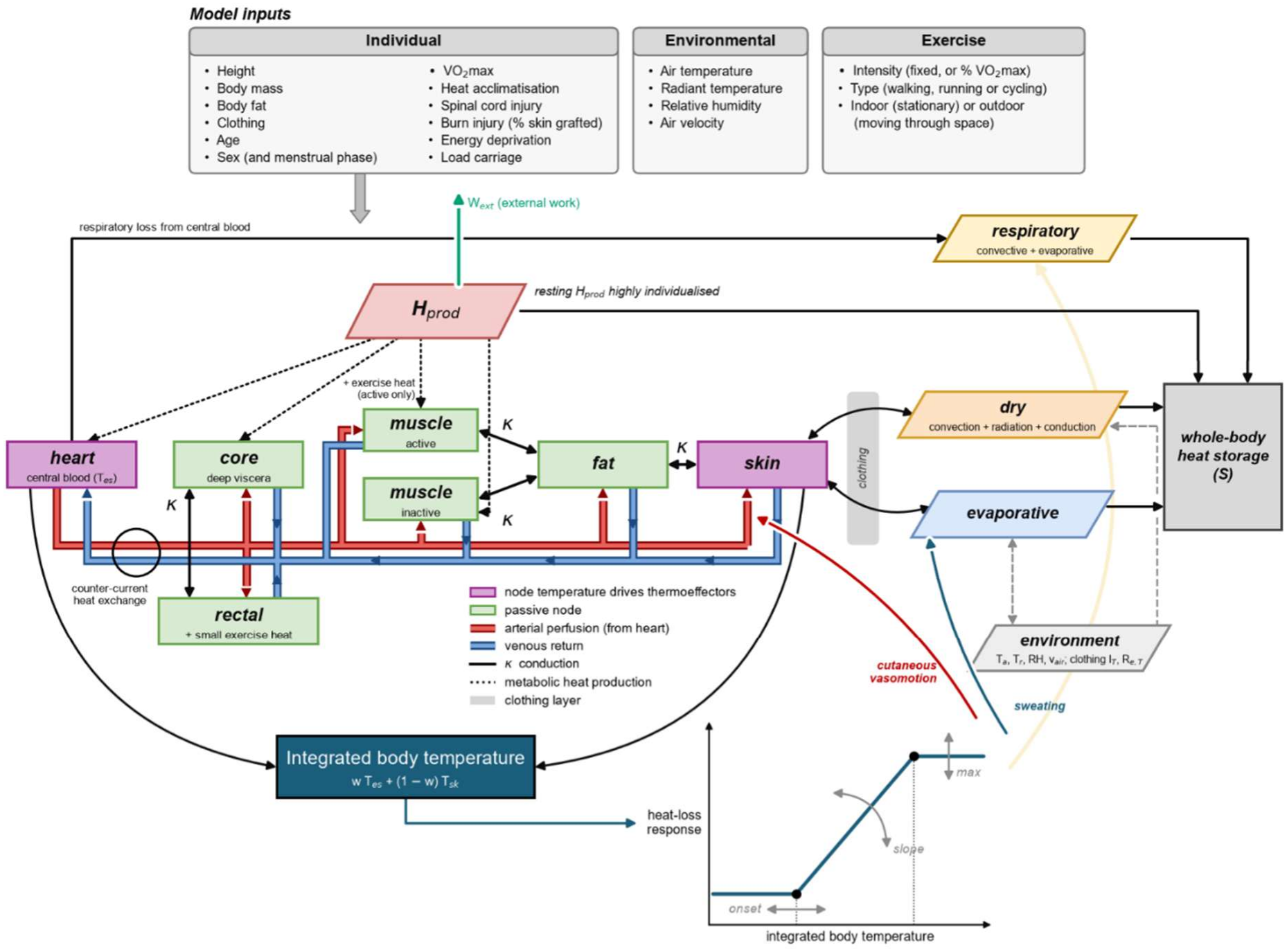
Schematic representation of TeMP’s input parameters, physical architecture and physiological control system.

We intentionally built TeMP’s passive system with low spatial resolution (relative to JOS-3 and Fiala) which renders TeMP easier to interpret, validate, individualise, and debug. The active system dictates the thermoeffector response and uses *T*_body_ as its primary input parameter, integrated from heart (*T*_heart_) and skin temperature (*T*_skin_). Specifically, the absolute *T*_body_ scales the heat-loss thermoeffector responses, with each thermoeffector (skin blood flow, sweating, ventilation) being activated at distinct but individualised thresholds (see ‘***Modelling individual thermoregulatory control’***). We chose *T*_heart_ as the main physiological controller input to *T*_body_ because it most closely represents central blood temperature, which is considered the thermal signal most tightly coupled to thermoeffector activation (Taylor *et al*., 2014). In contrast, rectal temperature (*T*_rectal_) lags considerably behind changes in *T*_heart_ and is a poor representation of thermoeffector onset. However, we still model *T*_rectal_ because it has stronger clinical utility.

A unique but powerful aspect of TeMP, building from Werner (2010), is that the model is decentralised. Specifically, it contains no ‘reference’ *T*_body_ that the prevailing *T*_body_ is compared against (see section ***‘no set-point architecture’***). In most thermophysiological models, the *error* between the reference and prevailing *T*_body_ determines the magnitude of the thermoeffector response. However, this concept is inconsistent with accepted physiology (Romanovsky, 2018), introduces unnecessary computational steps, and severely limits the scope for individualisation. In TeMP, the framework presented allows absolute tissue temperatures to emerge naturally in response to an individual’s own heat balance and individual characteristics.

### No set-point architecture

Numerous models operate based on the existence of a thermoregulatory “set-point”, defined as a fixed reference temperature from which a deviation (error) is computed, with that error driving effector output (Stolwijk & Hardy, 1966; Fiala *et al*., 2001; Kobayashi & Tanabe, 2013). These models impose a baseline *T*_body_, and effector functions are driven by Δ*T*_body_ from that fixed baseline. However, comparing *T*_body_ to some ‘reference’ would require a single site at which all afferent thermal signals are integrated to drive efferent thermoeffector responses, and the known circuitry does not support such a site (Ran *et al*., 2016; Romanovsky, 2018).

Thermoregulation is best described as a set of independent thermoeffector loops, each driven by its own combination of afferent inputs. TeMP applies this by assigning each thermoeffector a unique *onset temperature T*_body_ at which the feedback loop engages (Figure 6). Adopting this architecture is critical because baseline temperature itself varies substantially between individuals, driven largely by metabolic rate, clothing, and environment. Healthy older adults report resting core temperatures up to ~0.5 °C below young adults (McKenna *et al*., 2023) and patients with severely low metabolic rates, such as in anorexia, report near-hypothermic resting values (Wakeling & Russell, 1970). Modelling thermoeffector responses based on Δ*T*_body_ in these cases would cause heat loss to be activated at very low absolute *T*_body_, overestimating thermoeffector sensitivity in groups starting with a low baseline *T*_body_. Basal *T*_body_ can also be influenced by modulation of the thermoeffector loops themselves. For example, fever has been shown to increase the *T*_body_ onset threshold for heat-defence functions (Romanovsky, 2018). By removing the concept of a reference temperature and instead modelling baseline conditions using the framework shown in Figure 4, TeMP allows for more intuitive customisation of the thermoregulatory system according to personal and contextual factors.

**Figure 5.**
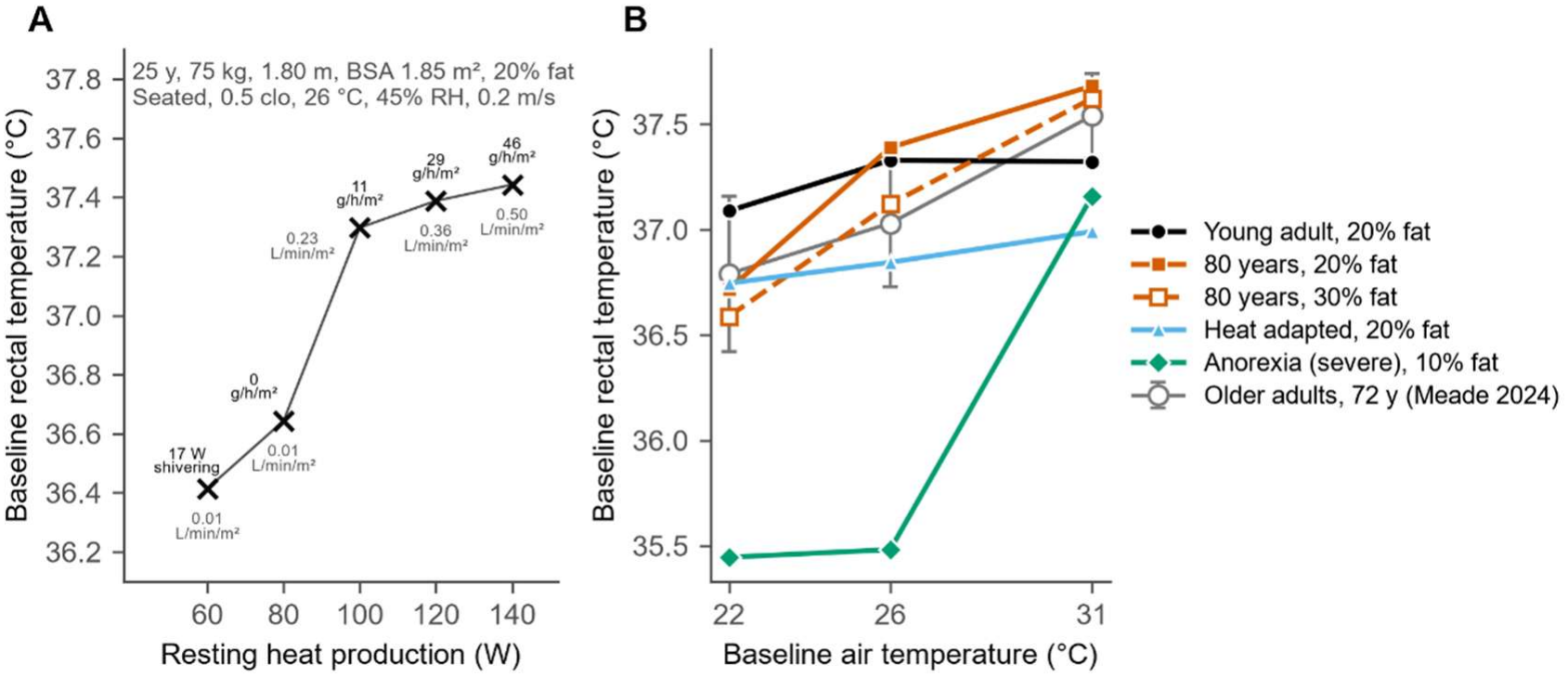
Resting simulation showing baseline rectal temperature and its primary modifiers. In a reference condition, panel A shows the direct impact of resting heat production on the prevailing steady state rectal temperature in a reference subject, assuming all else kept equal. Numbers above and below each datapoint are whole body sweat rate and skin blood flow, respectively. Panel B shows how individual modifications to resting metabolic heat production and vasomotor tone (latter relevant in older adults) impact baseline rectal temperature. In older adults, we also show the impact of fat-free-mass by showing an 80-year-old subject (all else kept equal) with a body fat of 20 and 30%, respectively. Comparative empirical data in older adults are also shown from Meade *et al* (2024a), however this study was not used to calibrate any model parameters.

**Figure 6.**
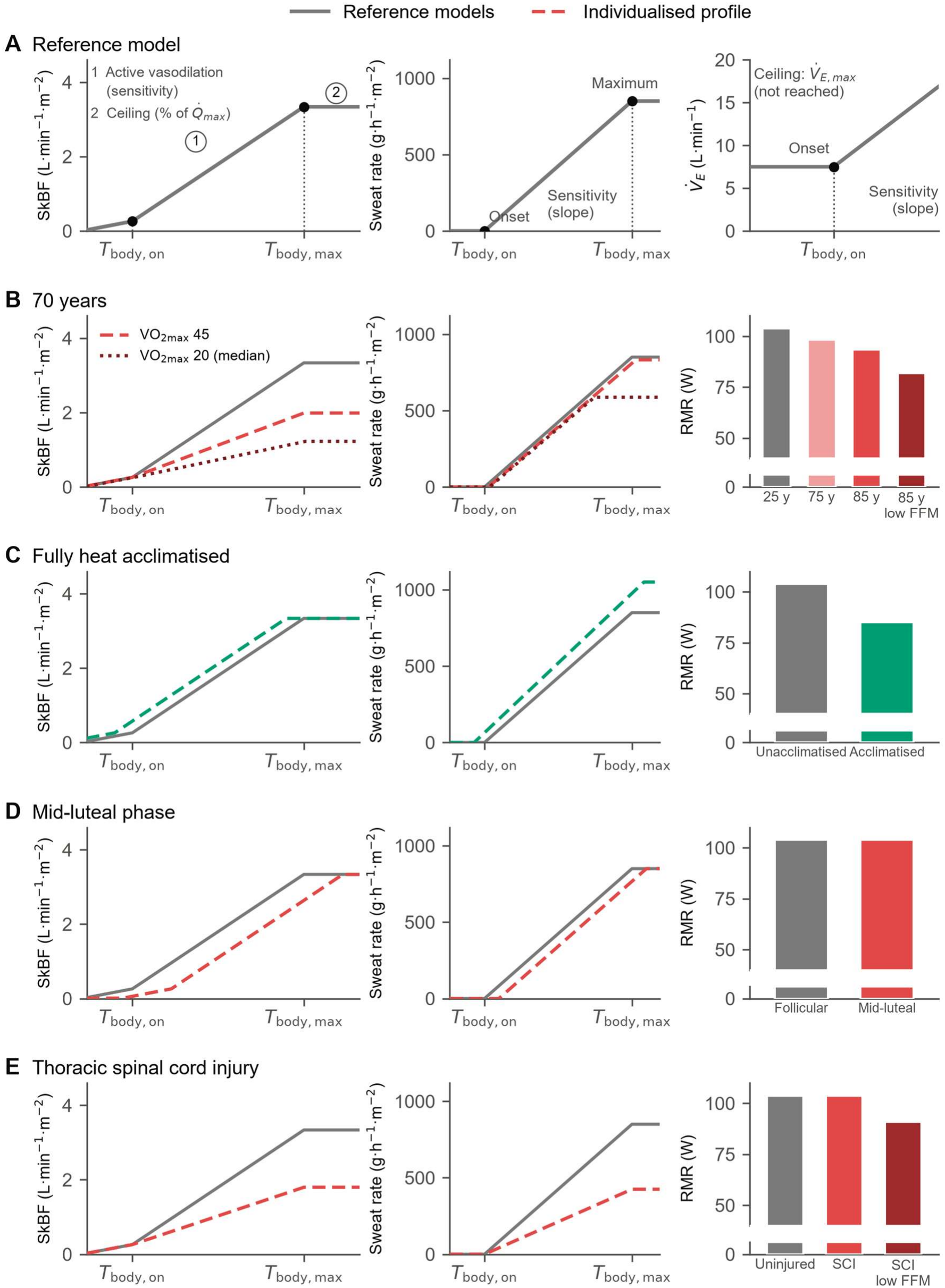
Simple piecewise equations linking mean body temperature to a given thermoeffector output and resting metabolic rate. Each thermoeffector function has an independent activation threshold (*T*_body,on_) and upper ceiling (*T*_body,max_). Skin blood flow (SkBF) has two inflection points which, from left to right, dictate i) the threshold for ‘release’ of vasoconstriction, and ii) cutaneous reflex vasodilation. The functions are designed to be consistent with known physiology and highly modular, allowing modification based on a multitude of individual factors (age, acclimatisation, spinal cord injury etc). Supplementary Table S2 provides the background evidence or logic supporting individual modifications.

### Baseline core and skin temperature prediction

Unlike extant thermoregulation models, baseline temperature in TeMP is not prescribed. Before each experimental condition, TeMP is run to steady state in a user-defined resting condition (environment, clothing and *H*_prod_) with all physiological effector loops active. The resulting node temperatures, i.e. the *T*_body_ at which heat loss matches heat production, become the starting state for the experimental simulation. This differs fundamentally from existing multi-node models which derive reference temperatures from a passive simulation (assuming thermoregulation ‘disabled’) in a predefined neutral environment and then defend them as set points (Fiala *et al*., 2001; Takahashi *et al*., 2021). In TeMP there is no reference temperature, and instead, baseline temperature is the closed-loop balance point between an individual’s heat production, body composition and effector characteristics (Werner, 2010).

Individual differences shift baseline temperature directly (Figure 5A and 5B). Resting *H*_prod_ is modelled across the lifespan (8 days to 95 years) from age and fat-free mass (Pontzer *et al*., 2021), and further modified for conditions such as heat acclimation (Lee & Kim, 2018) and anorexia nervosa (Wakeling & Russell, 1970). Resting *H*_prod_ from Pontzer is multiplied by 1.3 (i.e., 1.3 METS) to account for arousal and postural stability (Ainsworth *et al*., 2011). In line with standard heat balance concepts, a lower *H*_prod_ reduces the heat that must be lost, so a lower *T*_body_ is sufficient to achieve balance if thermoregulatory function is preserved. Hence baseline *T*_rectal_ falls with age, driven mainly by the age-related decline in *H*_prod_ and fat-free mass, and the clinically low *H*_prod_ of anorexia nervosa produces a correspondingly low resting *T*_body_. In anorexia, the lack of cold defence mechanisms at very low *T*_rectal_ is likely a torpor response driven in part by extremely low plasma leptin concentrations (Gavrilova *et al*., 1999). To account for this, we lower the *T*_body_ thresholds for shivering *H*_prod_ based on the ‘severity’ of energy deprivation but did not modify the heat-defence functions due to inconsistent data in these patients. Assuming all else is equal, Figure 5A demonstrates the impact of adjusting resting *H*_prod_ on baseline *T*_rectal_ in a reference subject in a comfortable environment.

### Modelling individual thermoregulatory control

Havenith (2001) sought to individualise effector thresholds, gains and upper limits by fitness, acclimatisation status and body composition, but within a set-point controller paradigm based on Gagge (1973). In TeMP, the control equations for whole body skin blood flow, eccrine sweating, and hyperventilation are governed by absolute *T*_heart_ and *T*_skin_, integrated to form *T*_body_. Consistent with empirical literature, the control of each function is governed by independent thermoeffector loops, with each exhibiting different onset thresholds, slopes, and max functions (Romanovsky, 2018). TeMP uses intentionally simple, modular, piecewise equations which link *T*_body_ to a given thermoeffector response (Figure 4). Empirical data supports the notion that one tissue temperature in isolation (i.e., core or skin only) is a poor predictor of thermoeffector activity. Instead, an appropriately weighted mean temperature of all tissues with thermal afferents predicts thermoeffector activity, expressed as an integrated *T*_body_ using:

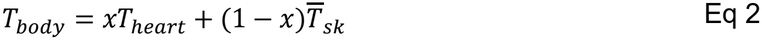

Where *T*_heart_ and 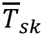 are absolute temperatures of the central circulation and *T*_skin_, respectively. The most appropriate weighting depends on the thermoeffector in question and is shown specifically for skin blood flow, eccrine sweating, and hyperventilation in their respective sections below.

As described in the below section ***‘Distinguishing between different ‘core’ temperature sites’***, TeMP can model the dynamic responses of different ‘core’ tissues, including the *T*_heart_ and *T*_rectal_. Here, we use *T*_heart_ as the ‘core’ site to determine the activation thresholds of different thermoeffectors, owing to its strong agreement with central arterial blood temperature (Shiraki *et al*., 1986) and the lagged rectal temperature response to heating (Au *et al*., 2019). Figure 6 shows the basic functions of key thermoeffectors relevant to heat exposure, including cutaneous vasomotion, eccrine sweating, and hyperthermic hyperventilation. The way each of these thermoeffectors are modelled and the data to support the approaches are discussed in more detail below.

While the precise threshold shifts need to be tuned and validated from a broader series of studies, Figure 6 demonstrates how different individual factors can be accounted for by adjusting the ‘reference’ piecewise equations. For example, the *T*_body_ thresholds for cutaneous vasodilation and eccrine sweating in older adults, and females in the mid-luteal phase of the menstrual cycle are shifted to the right, increasing the *T*_body_ required to achieve thermal balance. Conversely, heat acclimatisation shifts the onset thresholds for vasodilation and sweating to the left, resulting in a lower *T*_body_ to achieve thermal balance.

#### Distinguishing between different ‘core’ temperature sites

A single, homogeneous ‘core’ temperature does not exist in humans. In practice, values obtained instead depend on the tissue in which it is measured and on the exposure context, such as rest or PA (Shiraki *et al*., 1986; Mündel *et al*., 2016). In human experimental trials, core temperature is most frequently indexed by oesophageal, gastrointestinal or *T*_rectal_ (Taylor *et al*., 2014; Meade *et al*., 2025a). Each site differs in absolute values and the dynamic response to a given change in heat balance (Taylor *et al*., 2014). As noted above, we take oesophageal temperature as a proxy of *T*_heart_ as it is closely tracks central blood or heart temperature, and therefore the temperature of blood delivered to the brain(Cooper & Kenyon, 1957; Shiraki *et al*., 1986). Furthermore, in organs with limited heat exchange (such as the brain), arterial blood temperature is the main determinant of the organ temperature (Zaretsky *et al*., 2018). By contrast, *T*_rectal_ better reflects deep tissue temperature and can be more clinically relevant since heatstroke pathophysiology implicates damage to central organs, particularly the gastrointestinal tract, kidney, and liver (DeGroot *et al*., 2026). TeMP models the dynamic responses of different ‘core temperature’ measurement sites so it can be validated from a greater quantity of previous data, while allowing the user to focus on the most relevant tissue temperature for their context, i.e., thermoregulatory control or clinical relevance.

In TeMP, *T*_heart_ is the arterial source for every perfused compartment (Figure 1). The venous return from skin, muscle, fat, viscera and the extremities mix in this node, so each compartment exchanges heat with it in proportion to its blood flow and the arterial–tissue temperature difference. The node also loses heat through respiration and gains heat from local myocardial metabolism. The heart’s small heat capacity reproduces the heart’s fast response to passive and active heating (Cranston *et al*., 1954; Shiraki *et al*., 1986; Mündel *et al*., 2016; Au *et al*., 2019). At rest, cooler blood returning from the skin and muscle generally keep *T*_heart_ below *T*_rectal_. During PA, an initial ‘dip’ in *T*_heart_ is expected (Roberts & Wenger, 1979; Hirai *et al*., 1991) and is reproduced in TeMP (Figure 7). That brief reduction in *T*_heart_ is dependent on perfusion from cooler muscle into the heart, a transient behaviour that reverses as the muscle warms > *T*_heart_. The rectum, by contrast, is relatively well insulated, with low blood flow and low local *H*_prod_ (Taylor *et al*., 2014). In TeMP, the rectum gains heat by 1) perfusion from the heart node, which is low at rest and rises with PA intensity, 2) conduction from the lumped core compartment, and 3) via conduction from a small share of heat produced in the adjacent pelvic muscle during exercise. As a result, *T*_rectal_ lags behind *T*_heart_, responds slowly to heating (Au *et al*., 2019; Meade *et al*., 2024c), and equilibrates at a higher value (Mündel *et al*., 2016). Figure 7 shows the resulting *T*_heart_ and rectal responses of a young adult exposed to 40 °C and 10% RH while seated at rest or walking at 1.4 m·s^−1^.

**Figure 7.**
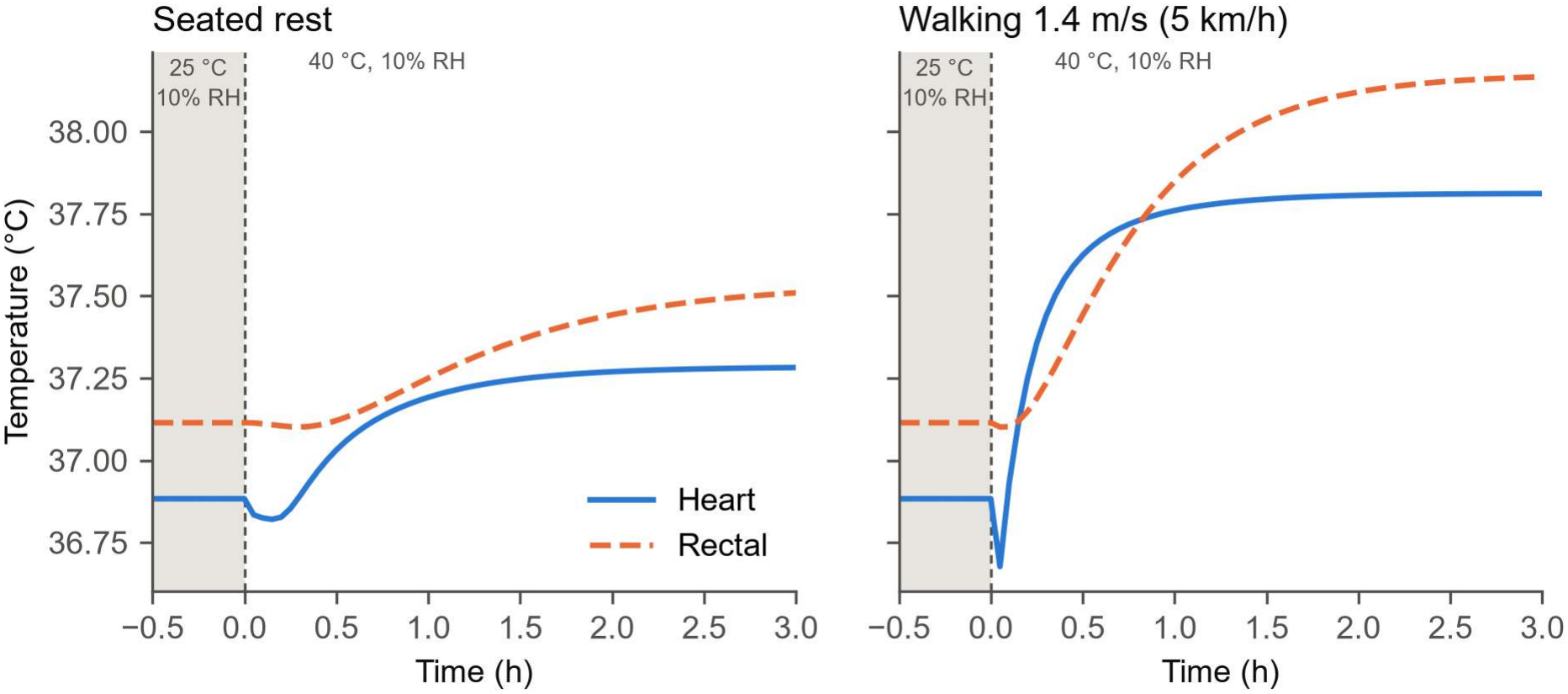
Dynamic heart and rectal temperature response to 40 °C, 10% RH at rest (grey box) and during physical activity in a simulated young healthy adult. Consistent with empirical data (Mündel *et al*., 2016; Meade *et al*., 2025b), heart temperature is below rectal temperature at baseline and rises faster during passive and physical activity heat stress. The equilibrium temperature is however greater in the rectum, as it is less impacted by venous return from cooler tissues. In the walking trial, the initial ‘dip’ in heart temperature is primarily explained by blood perfusion into cooler skeletal muscle returning to the heart.

#### Dynamic skin and muscle blood flow at rest and during physical activity

TeMP’s heart to skin blood flow functions deliver heat to the periphery where it can be liberated to the environment. We developed a novel algorithm to calculate skin blood flow building from earlier models (Fiala *et al*., 2001). Skin blood flow (SkBF) is constrained primarily by the limitations in maximal cardiac output (*CO*_max_) allocation, where there exists a ceiling on the fraction of cardiac output that can be directed to the skin, ensuring adequate perfusion to vital organs (Rowell, 1983). However, for a given *T*_body_, the fraction of *CO*_max_ directed to the skin decreases with progressive elevations in PA, while also shifting the *T*_body_ activation threshold to the right (Rowell, 1974; Taylor *et al*., 1988). Figure 8A shows that this response is minimal at low intensity PA but drops SkBF considerably as PA intensity reaches 60% We intentionally built TeMP’s passive system 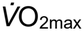 and above. Drawing from data where the *T*_body_ induced change in whole-body SkBF was estimated (Minson *et al*., 1998), we determined that in young healthy adults, approximately 24% of maximum cardiac output (*CO*_max_) is directed to the skin at the limit of thermal tolerance, corresponding to a *T*_body_ of ~38.8 °C (Minson *et al*., 1998). In older adults, this approach alone overestimates total SkBF (correcting for the reported 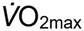 in older people), suggesting that age has an independent effect on maximal SkBF, at least in those aged >65 years. Such a view is supported empirically (Holowatz & Kenney, 2010), providing the rationale for including an age-related decline in SkBF on top of that induced by lowering individual fitness as shown in Figure 8B.

**Figure 8.**
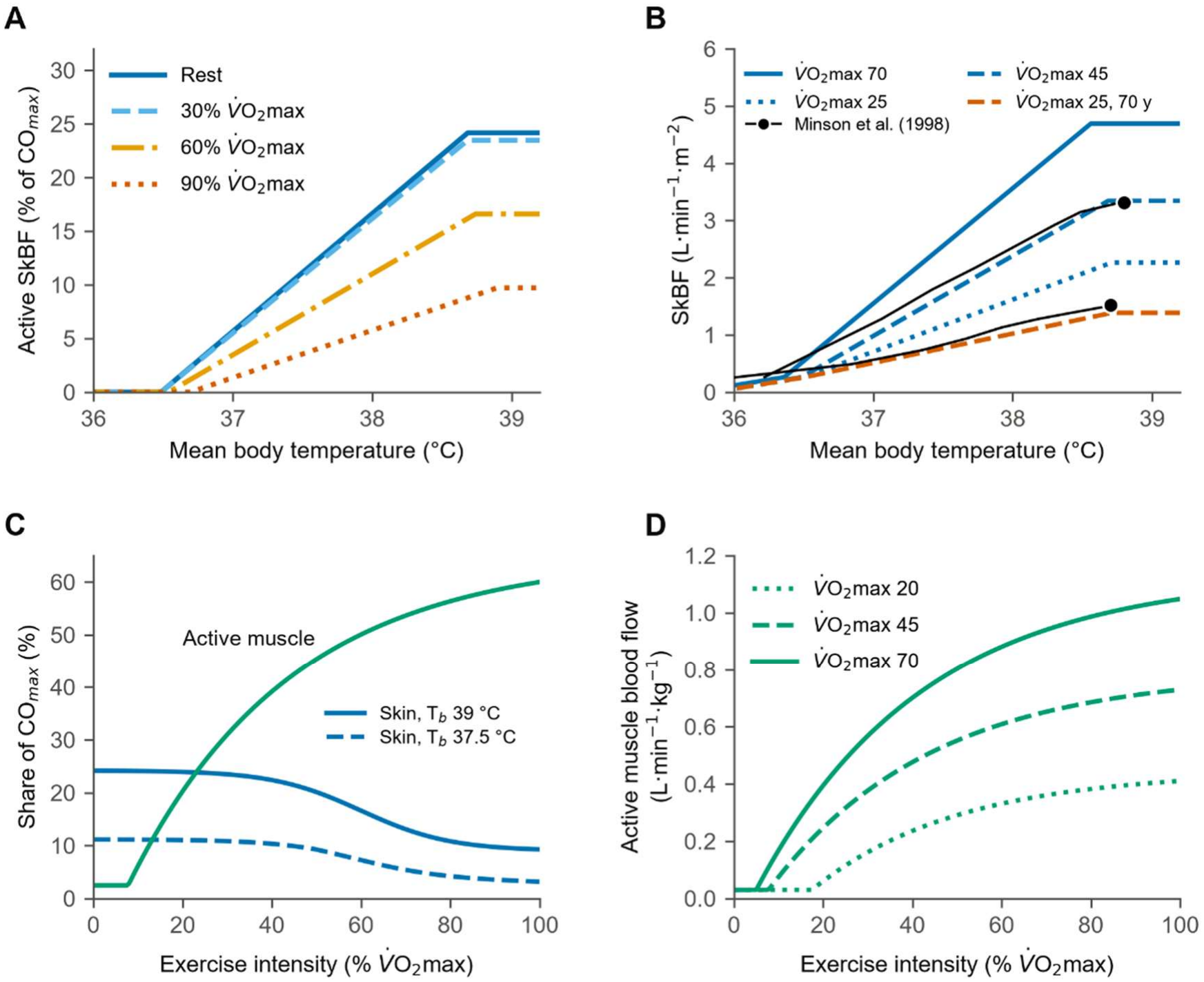
Graphical illustration of the dynamic skin (SkBF) and muscle blood flow (MBF) functions. SkBF is calculated as a fraction of maximum cardiac output allocation (*CO*_max_) which decreases based on the relative exercise intensity and also shifts to the right (A) consistent with (Rowell, 1974; Taylor *et al*., 1988). Constraining SkBF as a fraction of maximum cardiac output allocation (*CO*_max_) produces sensible flows that are neatly constrained by an individual’s maximum oxygen consumption. Panel B demonstrates how exercise and individual 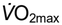 determines the absolute skin blood flow, with solid, dashed, and dotted lines showing an example for an individual with a 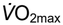 of 70, 45, and 25 ml/kg/min, respectively. The orange dashed line shows the response in an older participant with a 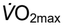 25 ml/kg/min. The fraction of *CO*_max_ reserved for active MBF is not impacted by body temperature and is instead a direct function of relative exercise intensity (C). Absolute active muscle blood flow is dependent on both individual 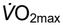 and relative exercise intensity and is based on data by Mortensen *et al* (2005) (D).

Individual 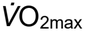 is used to estimate *CO*_max_ using the linear function below, assuming a linear relation between CO and 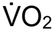 consistent with coefficients measured during incremental exercise and in earlier invasive studies (Åstrand *et al*., 1964; Beck *et al*., 2006).

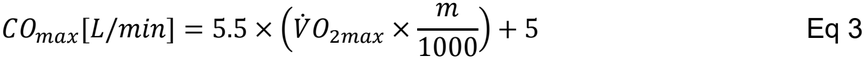

Where 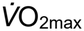 is maximal oxygen consumption in ml/kg/min, and m is mass in kg. Calibrated using data from Minson *et al* (1998), we set an upper ceiling for SkBF of 24% CO_max_. Using this approach results in a fully constrained level of SkBF regardless of individual 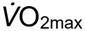 At 70 years, maximum skin blood flow is 44% lower than in a 25-year-old of the same fitness, and because the temperature range over which vasodilation develops is unchanged, the slope of the response falls by the same proportion. At present, TeMP continues the age-related decline in SkBF beyond this, plateauing at 40% of the reference value from age 80 years onward. We extrapolated beyond known data to better capture the well documented elevation in heat-related mortality in those aged 80-90+ years (Fouillet *et al*., 2006), but the reader should be aware that such age-related effects are currently a reasonable best guess.

Active muscle blood flow (*MBF*_active_) is also bounded by *CO*_max_ and has its own unique allocation depending on the PA, expressed as % 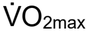 Based on suggestions by Rowell (1974), *CO* allocation to SkBF decreases as intensity increases (Figure 8C), likely secondary to increased skin sympathetic nerve activity at higher intensities (Taylor *et al*., 1988). Figure 8C shows maximum *CO* allocation to the skin and active muscle, and is based on theoretical models proposed by Rowell (1974).

Importantly, the *CO*_max_ model shown in Figure 8 is only relevant for SkBF above the active vasodilation threshold. Left of threshold, we assume absolute values based on *T*_body_ directly. While only active vasodilation is shown in Figure 8, across the entire range of *T*_body_, SkBF is modelled as a five-parameter piecewise function, reflecting its variable control for both heat and cold defence. Modifiable parameters include i) a minimum under maximal vasoconstriction, ii) an onset threshold for the release of vasoconstriction, iii) an onset threshold for active vasodilation, iv) a rising slope (gain) once vasodilation begins, and v) a maximum SkBF, capped by *CO*_max_ bounded cardiac output allocation as described above. This structure captures the two distinct neural pathways governing skin blood flow. First, the withdrawal of vasoconstrictor tone at low thermal load, followed by active vasodilator-mediated flow increases once a higher, separate threshold is exceeded. This approach is more mechanistically consistent than treating SkBF as a linear function of *T*_body_.

The control equations for skin blood flow use a *T*_heart_ to *T*_skin_ weighting ratio of 0.8:0.2. Passive release of vasoconstrictor tone (*SkBFp*) is modelled as:

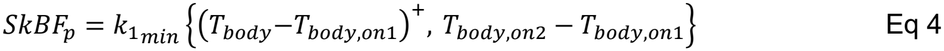

where *T*_*body*,on*1*_ and *T*_*body*,on*2*_ are the onsets of vasoconstrictor release and active vasodilation, respectively. *k*_1min_ (0.4 L·min^−1^·m^−2^·°C^−1^) is the slope of vasoconstrictor release. + implies that only a positive value can be used, if else 0. As in Figure 8A, the ceiling on maximal skin blood flow is modelled as:

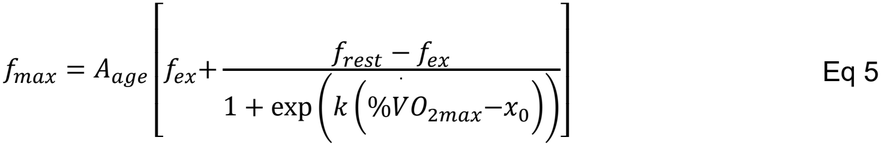

Where *A*_age_ is an age dependent attenuation on allocation of *CO*_max_ to the skin, *f*_rest_ = 0.24 and *f*_ex_ = 0.09 are fractions of *CO*_max_ available to skin at rest and at maximal PA, respectively. *k* (10) and *x*_0_ (0.6) show the steepness and midpoint of their transition, respectively. % 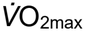 describes the relative PA intensity.

Active cutaneous vasodilation (*f*_active_) is modelled as:

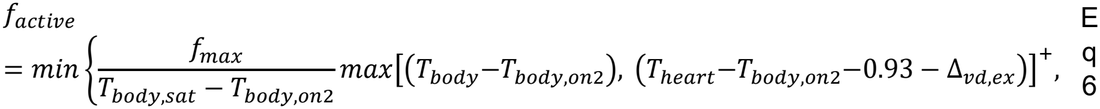

Where *T*_*body*,sat_ is the *T*_body_ where vasodilation saturates. *T*_heart_ can override inhibitory effects of cold skin on vasodilation if it is raised 0.93 above *T*_*body*,on2_. Δ_*vd,ex*_ is the rightward shift in *T*_body_ onset for vasodilation relative to the exercise intensity (Taylor *et al*., 1988):

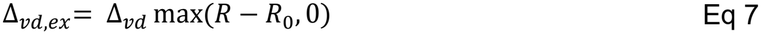

Where *R* is the fraction of 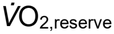 above resting, *R*_0_ = 0.45 and Δ_*vd*_ = 0.6 °C. Exercise at a 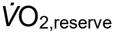 > *R*_0_ delays the onset of active vasodilation in proportion to intensity (Taylor *et al*., 1988). The shift is expressed in *T*_heart_, moving the *T*_body_ drive by 0.8 Δ_*vd,ex*_ and the *T*_heart_ override by the full amount. Saturation moves with the onset, so the slope *f*_max_/(*T*_*body*,sat_ – *T*_*body,on2*_) is unchanged, consistent with Taylor’s finding of no change in slope.

Active skin blood flow (L·m^-2^·min^−1^) is then modelled as:

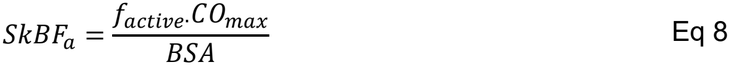

Where BSA is body surface area in m^−2^. The final term for overall skin blood flow becomes:

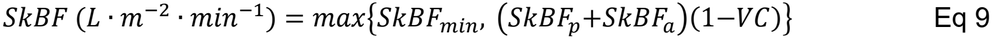

Where *VC* is the cold vasoconstrictor drive (0 in the heat). Threshold values are for the reference adult and shift with individual characteristics (supplementary Table S2).

#### Whole body sweating at rest and during physical activity

At rest, TeMP models whole body sweating based *T*_body_, weighted 0.9:0.1 for the *T*_heart_ and *T*_skin_, respectively. The current version adopts values of 0.01 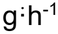·m^−2^ for baseline, a *T*_body_ onset threshold of 36.75°C and a slope of 164.4 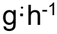 ·m^−2^·°C^−1^ fitted to passive environmental chamber data (Meade *et al*., 2023; Meade *et al*., 2025b), and a max of 800 ġ·h^−1^·m^−2^ based on Havenith (2001). These parameters form the ‘base’ model which can be adapted for individual characteristic such as fitness, age, menstrual cycle phase, skin grafting, and heat acclimatisation. Interestingly, this approach will almost always underpredict sweating during PA (de Korver *et al*., 2025) because additional thermal and non-thermal signals drive sweating before a measurable increase in *T*_body_ (Shibasaki *et al*., 2006; Notley *et al*., 2024). The ‘extra’ sweating during PA is at least partly explained by factors such as elevated muscle temperature, central command, and metaboreceptor activation, all of which can increase sudomotor sensitivity before *T*_body_ rises significantly (Kondo *et al*., 2000). Modelling each of these factors is beyond the current scope of TeMP, so we include an approach where the ‘reference’ sweating model becomes more sensitive as relative intensity progressively increases (Kondo *et al*., 2000). We scale intensity using 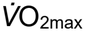 reserve (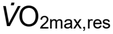) instead of % 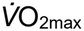 because low fitness individuals could operate ~15% 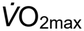 at rest.

In Figure 9, we show the utility of this model in predicting *T*_rectal_ responses to PA, calibrated against three empirical studies involving treadmill running (Che Muhamed *et al*., 2016) and cycle ergometry at a variety of relative intensities (Mora-Rodriguez *et al*., 2010; Périard *et al*., 2012). Comparisons against additional work are shown for observational purposes (Saltin & Hermansen, 1966; Drinkwater *et al*., 1976). Further, exercise metabolic cost rises 2.3% per °C of core temperature above 37 °C, representing the reduced work efficiency observed during heavy exercise in the heat (Fink *et al*., 1975). Future validation will include more studies to determine the accuracy of this approach in a range of different environments, PA modes, and relative intensities.

**Figure 9.**
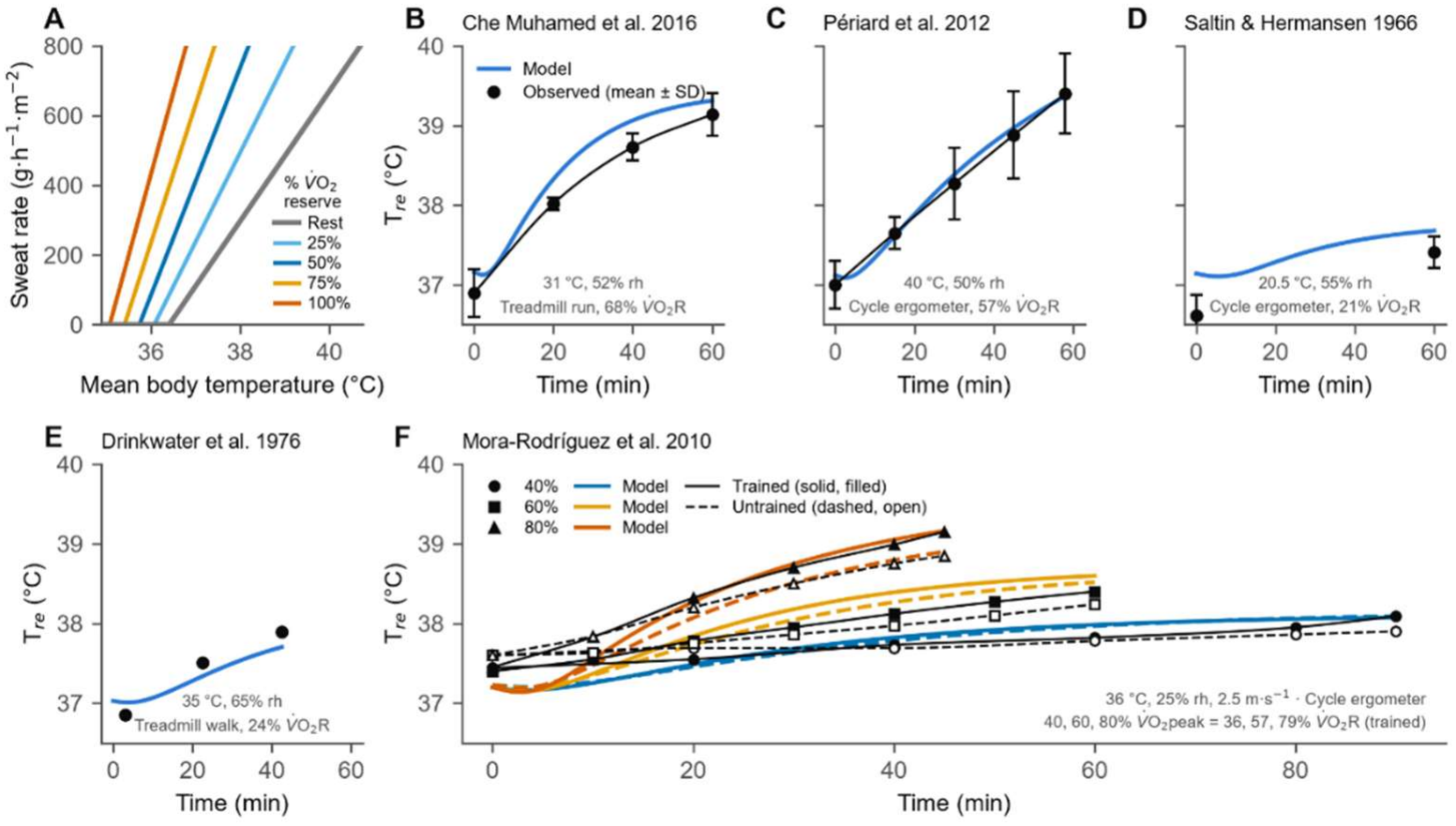
Visual representation of the physical activity sweating model. During physical activity, thermal and nonthermal factors activate eccrine sweating before measurable increases in body temperature. Panel A shows the additional sweating sensitivity relative physical activity intensity above resting (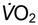 reserve). Panels B-F show the effect of the physical activity sweating model on TeMP’s emergent rectal temperature (blue, orange, and yellow lines) and compares those traces to that observed (black lines). Data from Drinkwater (1976) and Saltin & Hermansen (1966) were not used to calibrate the model.

The sweating function is driven entirely by *T*_heart_ and *T*_skin_, using a respective weighting of 0.9:0.1 represented as *T*_body,sw_. The precise function for thermal sweating is modelled according to Nadel *et al*. (1971), with PA modifying its sensitivity:

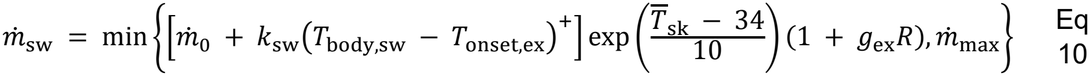

where 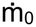 is the baseline sweat rate, *k*_sw_ the sweating gain (g·h^−1^·m^−2^·°C^−1^), 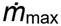 the individual’s maximum sweat rate (Eq 12), *R* the relative intensity (0–1, fraction of 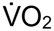 reserve), and *g*_ex_ = 1.5 is the fractional rise in sweating sensitivity at maximal intensity. During PA, the onset threshold also falls in proportion to relative intensity:

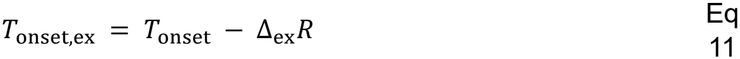

where *T*_onset_ is the individual’s resting onset threshold and Δ_ex_ = 1.5 °C is the fall in onset at maximal intensity. At rest (*R* = 0) both modifiers vanish and Eq 10 reduces to the thermal function only. 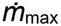 is unaffected, as maximal sweat rate reflects sweat gland capacity rather than acute PA. 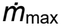 scales with fitness and acclimatisation (Havenith, 2001):

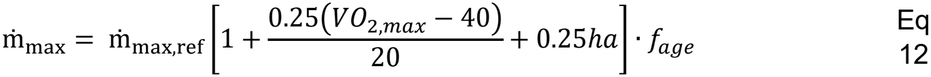

Where 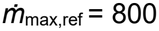 g·h^−1^·m^−2^ represents maximum sweat gland output for an unacclimatised adult with a 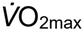 of 40 ml·kg^−1^·min^−1^. 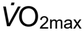 range is limited from 20-60 ml·kg^−1^·min^−1^, *ha* is heat acclimatisation state (0-1) and *f*_age_ is the age reduction in sweating capacity (Table S2).

#### Hyperthermic hyperventilation

Previous models include ventilatory heat loss by assuming a linear relationship between 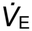 and metabolic rate (Fiala *et al*., 2001; Malchaire *et al*., 2001; Potter *et al*., 2017; Takahashi *et al*., 2021). In TeMP, we aim to be more physiologically consistent by considering that 1) above a threshold, *T*_body_ directly increases minute ventilation (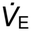) (Gibbons *et al*., 2022a; Gibbons *et al*., 2022b) and 2) a first ventilatory threshold (*V*_T1_), where 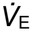 rises disproportionately to 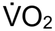 at higher relative PA intensities ~60%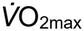 (Wasserman, 1987). At rest, modelling 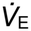 appropriately may have only a small impact on total heat loss, but the downstream effects on acid-base balance will become highly relevant in more detailed modelling extensions of the cardiorespiratory system. Further, during high intensity PA above *V*_T1_, the added heat loss is relevant for improving the accuracy of TeMP’s emergent temperatures. An example output of the model at result and during two exercise intensities is shown in Figure 10. The maximum 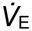 from an individual subject is predicted from individual 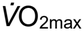 (Blackie *et al*., 1991):

**Figure 10.**
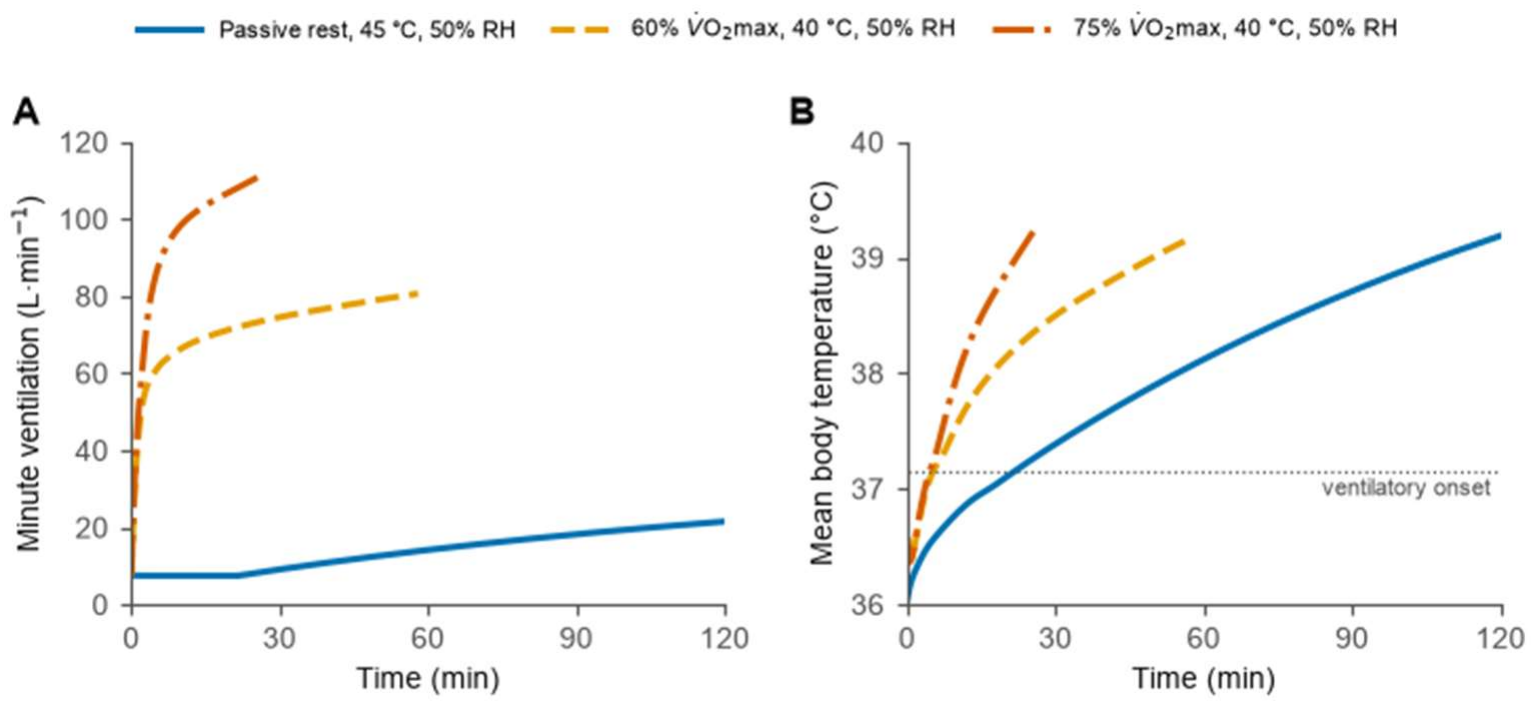
Ventilatory response to passive and active heat stress. Panel A shows a passive uncompensable heat exposure (solid blue line), and cycling exercise at a relative intensity of 60 (dashed yellow line) and 75% (dashed and dotted orange line) 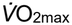, respectively. Panel B shows the mean body temperature trace throughout each exposure, with the horizontal dotted line showing the onset of hyperthermic ventilation.

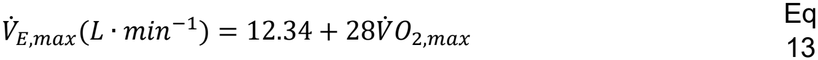

Where 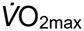 is in L·min^−1^. To capture the effects of PA intensity above *V*_T_ and *T*_body_, we first calculate 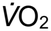 as:

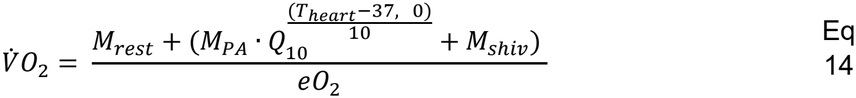

Where *M*_rest_, *M*_PA_, and *M*_shiv_ are resting, PA, and shivering metabolic rate in Watts. Accounting for a *Q*_10_ effect (1.25), *M*_PA_ is elevated by *Q*_10_ once *T*_heart_ rises >37 °C during exercise. *eO*_*2*_ is the energy equivalent of O_2_ and is equal to 348.3 Watts per L·min^−1^.

To account for PA intensity above *V*_T1_, ventilatory equivalent (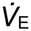/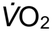) is modelled as:

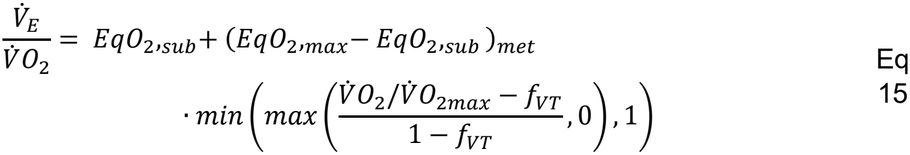

Where *EqO*_*2*_,_sub_ and *EqO*_*2*_,_max_ are the ventilatory equivalents below *V*_T1_ (25) and at 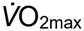 (35), respectively (Norton *et al*., 1995). *f*_VT_ is the fraction of relative 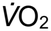 demarking the *V*_T_ (0.6). Total 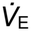 considering 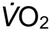, ventilatory equivalent and *T*_body_ is then:

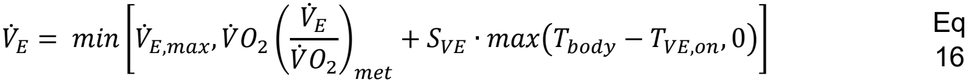

Where 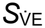is the slope for the rise in 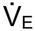with *T*_body_ (6.9 L·min^−1^·°C^−1^) with an onset *T*_body_ of 37.15 °C, each based on Gibbons *et al* (2022a; 2022b).

Respiratory convective heat loss (*C*_res_) is calculated from:

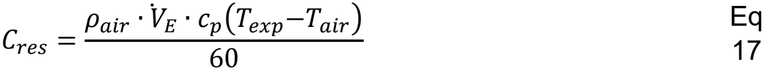

Where *ρ*_air_ is air density (kg·L^−1^) and *c*_p_ is the specific heat of moist air (J·kg^−1^·°C^−1^). *T*_air_ is the air

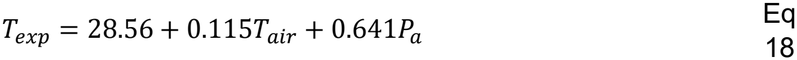

Where *P*_a_ is ambient water vapour pressure (kPa). Respiratory evaporative heat loss (*E*_res_) is calculated from:

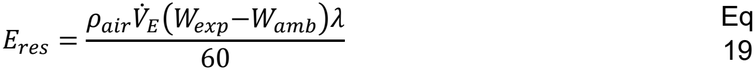

Where *W*_exp_ and *W*_amb_ are the humidity/mixing ratios of expired air (saturated at *T*_exp_) and ambient air in g·kg^−1^, respectively. λ = 2430 J·g^−1^.

### TeMP’s dynamic responses to compensable and uncompensable heat stress

Predicting individualised tissue temperature trajectories over sustained heat exposure is highly relevant for public and occupational health, especially with the growing impact of climate change. Figure 11 shows a model output for a compensable (40 °C, 10% RH) and uncompensable (40 °C, 70% RH) condition in a variety of subject types. We do however emphasise that an environment being ‘compensable’ does not necessarily mean it is safe for prolonged exposure, as the operating point for *T*_body_ may achieve thermal balance, but invoke unsustainable physiological stress (Foster *et al*., 2025). Before the experiment, dry heat loss would have been sufficient to achieve *S* ~ 0. Note however that in the experimental conditions, due to vasodilation, *T*_skin_ begins to increase, and with it reducing the environment to *T*_skin_ temperature gradient. However, the rise in *T*_skin_ has physiological and biophysical limitations, such that vasodilation alone cannot fully restore heat balance in hot environments. For the duration that *S* is positive, *T*_body_ will continue to rise.

**Figure 11.**
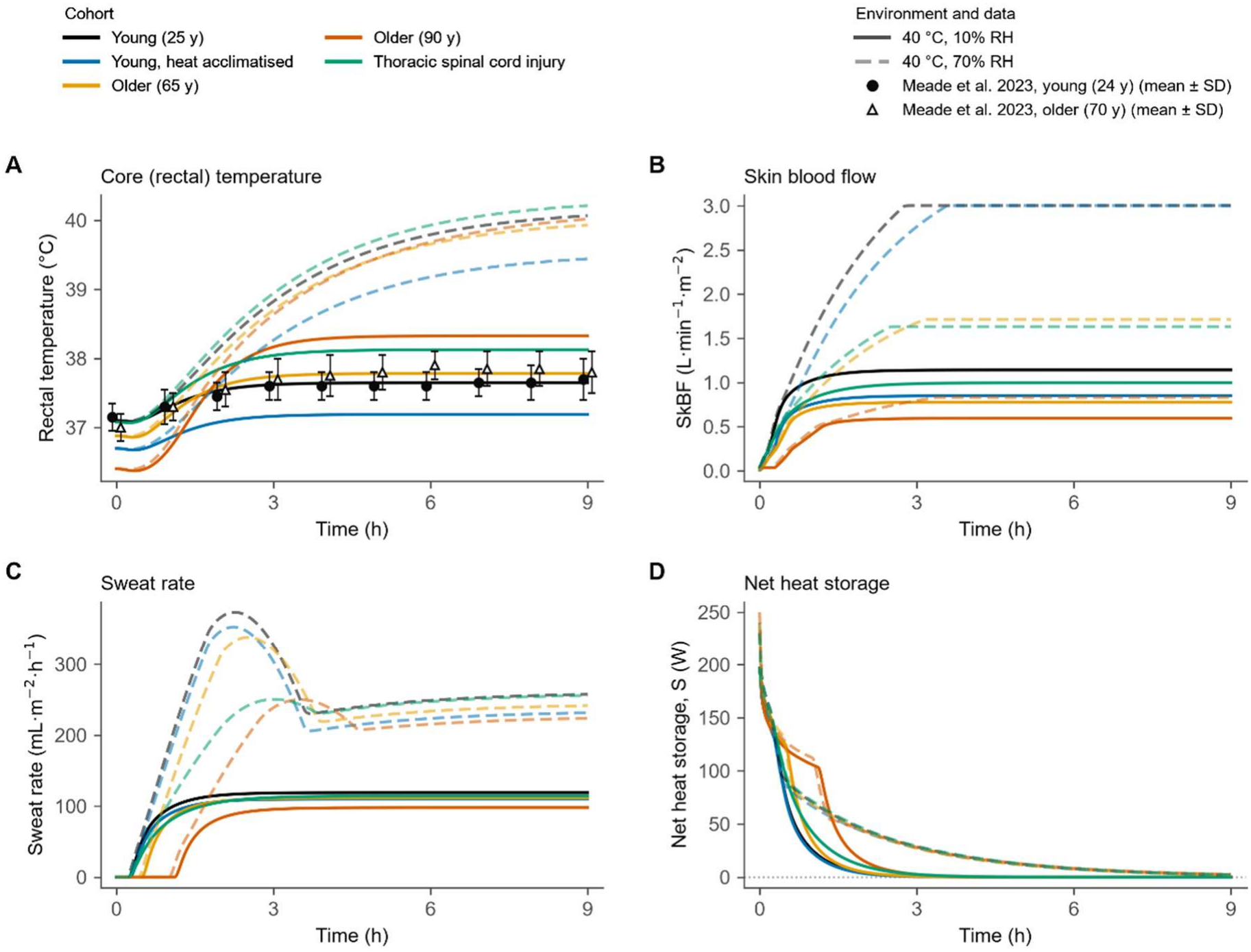
Model output for multiple participant types during a compensable (solid lines, 40 °C, 10% RH) and uncompensable (dashed lines, 40 °C, 70% RH) passive heat exposure. Note the large differences in absolute rectal temperature in the 10% RH condition between participants, highlighting the benefits of heat adaptation and the risks associated with ageing and spinal cord injury. In line with the underlying philosophy of TeMP, the net heat storage traces are similar across participants, but the body temperature *required* to elicit necessary heat loss differs dramatically between groups. Symbols represent published data in the same condition in young (filled circles) and older adults (open triangles), of which data were used to calibrate the ageing effect and rectal temperature trajectory (Meade *et al*., 2023). In the 70% RH condition, sweat evaporation is inefficient in all participants, but note how the heat acclimatised individual still operates at a lower rectal temperature throughout. The observed effect occurs secondary to a more rapid heat loss response and lower heat production. In the 70% RH condition, also note the transient decrease in sweat output (panel C dashed lines) which occurs secondary to hidromeiosis (Candas *et al*., 1983), a well-documented phenomenon which decrease sweat drippage from already saturated skin.

Once *T*_body_ reaches its onset threshold, eccrine sweating begins and increases the rate of evaporative heat loss. Figure 10 demonstrates that in both the compensable and uncompensable situation, evaporative heat loss is activated ~5 to 40 minutes into the exposure dependent on the subject’s individual thermoregulatory control (see Figure 6). Since the driver for thermoeffector functions is *T*_body_, it will itself plateau if a state of thermal equilibrium can be established. However, if an intrinsic factor impairs some thermoeffector function, the lever (*T*_body_) that drives heat loss will simply equilibrate at a higher temperature. In the low humidity condition (solid lines), this effect is shown clearly in the simulated individual with a thoracic spinal cord injury and older adults, who require a higher *T*_body_ to drive necessary heat loss. In the 70%RH condition (dotted lines), the heat loss achievable through sweat evaporation *cannot* drive S back to 0 within safe *T*_body_ limits. Here, we show that TeMP can define the environmental boundaries whereby *T*_body_ cannot be regulated, and the individual factors determining the proximity to those boundaries.

### Comparison with existing models (high level summary)

Supplementary Table S1 provides specific differences between TeMP and a variety of commonly used thermal models. TeMP represents a fundamental architectural departure, using *absolute* integrated *T*_body_ to drive independent, effector-specific feedback loops with unique onset thresholds, slopes, and physiological ceilings. TeMP’s decentralised control architecture aligns closely with modern physiological consensus regarding thermal afferent integration (Meade *et al*., 2024b), built from foundations laid by earlier works introducing these concepts (Werner, 2010). Fiala’s controller was fitted to responses of young adult males and, to our knowledge, has not been re-derived for other populations. JOS-3 adopts regional multipliers on sweating and vasodilation, applied as a step from age 60 y, without altering effector thresholds (Takahashi *et al*., 2021). Further, JOS-3 uses a ‘bottom-up’ approach to calculate *CO* from the sum of all regional flows, meaning that flow is not constrained by individual fitness levels. In contrast, TeMP uses a ‘top-down’ approach where flow is bounded by *CO*_*max*_, bounding skin and muscle blood flow from individual fitness and a realistic *CO* allocation.

Simpler frameworks like PHS, HSDA, and HEAT-Lim do not simulate thermoregulatory control. HEAT-Lim is a single compartment heat balance model that becomes relevant only when heat storage is greater than zero (uncompensable), relying on fixed *T*_skin_ and *H*_prod_ to calculate core temperature changes from heat storage alone (Vanos *et al*., 2023). PHS (Malchaire *et al*., 2001) and HSDA (Potter *et al*., 2017) are largely empirical models that predict core temperature for a standard (non-individualised) subject from user-specified conditions. In contrast, TeMP’s tissue temperatures emerge naturally from an individual’s unique profile, thermoregulatory function and heat balance, allowing it to accommodate a wide profile of individual differences without widespread re-fitting of physiological parameters.

## Limitations

There are several limitations that should be considered when interpreting the model. Firstly, maximum values for SkBF and sweating are not taken from data because plateaus in each of these responses have not been observed empirically. Obtaining such data in resting participants is challenging as the thermal drive required would likely result in heatstroke. Secondly, TeMP does not yet incorporate a dynamic ‘dehydration’ effect, which during prolonged heat exposure would progressively decrease SkBF and sweating rate. Thirdly, the maximal SkBF responses during hyperthermia and exercise are not well characterised empirically due to the invasive nature of those measurements. Therefore, the skin and muscle blood flow allocations during passive and active heat exposure are anchored from a single study and a theoretical *CO* allocation to muscle at different exercise intensities (Rowell, 1974; Minson *et al*., 1998). Finally, the downstream cardiorespiratory effects are not considered, but we are actively integrating TeMP with a dynamic cardiopulmonary model to predict downstream effects of hyperthermia and progressive fluid loss on heart rate, regional blood flow, and acid-base balance (Whittle & Diaz-Artiles, 2021).

## Future directions and conclusions

The framework presented lays the foundation for parameter tuning and validation protocols based on vast numbers of previous studies in diverse contexts and individual factors. In addition, follow up manuscripts will detail the cold defence functions and integration with a detailed cardiopulmonary model (Whittle & Diaz-Artiles, 2021). While it is unknown how many health conditions and medications impact thermoregulatory function, TeMP now provides immediate justification for physiologists to generate data that can be incorporated into the model. With greater individualisation coupled with an integrated cardiopulmonary model, TeMP has the potential to be adopted as tool to aid in heat-risk forecasting, decision making, and education delivery.

## Acknowledgements

JF expresses sincere gratitude to the late Professor George Havenith for sparking his interest in thermoregulation modelling and discussions related to individualisation approaches. JF would also like to thank Oliva Simons for their thoughtful discussions and input relating energy deprivation to baseline hypothermia, which helped anchor the link between metabolic rate and baseline temperatures.

## Author contribution

JF and RDM conceived the original model architecture. JF produced the code and was the primary developer, with consistent feedback from RDM during its development. RW set up the code structure in Julia using *ModelingToolkit* and provided engineering assistance to JF throughout. Model code changes were made based on comments from GPK, CGC, SRN, AD, PDH, and SL at various stages. All authors provided feedback on the manuscript contributed to the text. All approved the final version of the manuscript.

## Code availability

TeMP is proprietary software and its source code is not publicly available. The model structure, governing equations and principal parameter values are provided in the manuscript and Supplementary Table S2. The code may be made freely available for non-commercial research purposes under a licence agreement, on reasonable request to the corresponding author.

## AI declaration

The lead author (JF) used an AI coding model (Anthropic, Claude Opus) to assist with implementing and debugging parts of the model code. The AI coding model was also used to compare TeMP’s dynamic outputs with previous literature used for model calibration and figure generation. All of TeMP’s parameter choices and equations were developed by JF with input from coauthors. The broad model architecture, written in Julia, was developed by RW.

