## Supplementary material for "A new thermophysiological model unifying heat balance theory and physiological feedback control: *Framework, heat defence functions, and individualisation*"

**Table S1.** Comparison of TeMP with selected thermophysiological models.

| | TeMP | Fiala et al. (2001) | JOS-3 (2021) | HEAT-Lim (Vanos et al. 2023) | Gagge two-node | PHS (ISO 7933) / HSDA |
|---|---|---|---|---|---|---|
| ***Effector control signal*** | Absolute integrated body temperature ($T_{heart}$ + $T_{skin}$, weighted per effector); independent threshold, gain and ceiling per effector | Error from head-core (hypothalamic) and mean skin set points, with non-linear (tanh) gains regression-fitted to 26 experiments; rate of skin cooling adds to cold responses | Error from head-core and segmental skin set points, with fixed linear gains (Stolwijk-based) and age decrement factors | No dynamic control: heat balance in which evaporation is limited by ambient humidity, maximum skin wettedness and maximum sweat rate | Error from core (36.8 °C) and mean skin (33.7 °C) set points; skin blood flow, sweating and shivering driven by warm and cold error signals with fixed gains | No feedback control. PHS: required sweat rate, capped; empirical skin temperature; core rises 0.0036 °C per W·m$^{-2}$ above 55. HSDA: Givoni–Goldman equilibrium core temperature (raised as $E_{req}$ exceeds $E_{max}$), approached exponentially |
| ***Baseline (resting) temperature*** | Emerges from the individual's own heat balance at rest; effector thresholds are independent of it | Set points = passive steady state of the reference body, nude, at 30 °C | Set points = passive steady state of the individual, nude, at PMV-neutral conditions; authors note insufficient data to support them | Fixed starting "core" temperature (37.0 °C, normothermic) at the start of each exposure window | Fixed set points (core 36.8, skin 33.7 °C) | Fixed starting values (PHS: rectal and core 36.8 °C, skin 34.1 °C; HSDA defaults: core 36.8 °C, skin 33 °C) |
| ***Adding individual factors*** | Modular: change only the parameters the evidence supports (thresholds, gains, ceilings, resting metabolism); baseline and responses then emerge from heat balance, with no set points to re-derive | Control equations regression-fitted to young, unacclimatised subjects; a new population would require refitting | Age "decrement signals" are fixed regional multipliers on effector output (sweating 0.40–0.75 and vasodilation 0.31–0.91, applied as a step at ≥ 60 y; shivering in decade steps), leaving thresholds unchanged; set points re-derived from a steady-state run of the individual | Fixed profiles defined primarily by maximum sweat rate and skin wettedness | Not supported beyond body size; new populations would need new set points and gains | Empirical corrections only: acclimatisation (both) and dehydration (HSDA). ISO 7933 does not predict "individual subjects"; HSDA "does not provide individualized predictions" (Potter et al. 2017) |
| ***Resting metabolic rate*** | Pontzer (age, fat-free mass) × 1.3 for seated rest; acclimatisation and energy-deprivation modifiers | Reference-person value (87.1 W), with local Q10 effect | Harris–Benedict (height, mass, age, sex); body composition not considered | 1.5 METs (1.8 W·kg$^{-1}$) assumed | User input | User input |
| ***Metabolic rate during activity*** | Individual resting rate + ACSM activity increment (walking, running; load scales by (m + L)/m); cycling from external power; cost rises with core temperature | User input | Physical activity ratio (PAR) multiplying basal metabolism | Rest (1.5 METs) for survivability; for liveability, the maximum sustainable metabolic rate is the output | User input | PHS: user input. HSDA: user input or the Pandolf load-carriage equation (load, speed, grade, terrain) |
| ***Skin blood flow*** | Ceiling is a fraction of maximal cardiac output (from $\dot{V}O_2$max), falling with relative exercise intensity; vasodilation onset delayed at high intensity | Fixed maximum, reduced linearly as muscle blood flow rises with exercise; not scaled to fitness | No maximum; multiplicative signal on segmental basal flow (scaled by cardiac index); cardiac output is the sum of flows, not a limit | Not modelled (no skin blood flow). $T_{skin}$ fixed at 35 °C | Driven by core and skin error signals; fixed maximum (90 L·m$^{-2}$·h$^{-1}$); not scaled to fitness | Not modelled |
| ***Exercise (non-thermal) sweating*** | Sweating onset threshold and gain modified by %$\dot{V}O_2$ reserve | Thermal error signals only; authors note poorer sweat prediction in heavy exercise, attributed to non-thermal factors | None; limb work heat enters segment core nodes (only head and pelvis have muscle layers); validated at rest only | Considered only through elevations in metabolic rate (interacting parameters, e.g. maximum sweat rate, must be tuned manually) | None; work heat is added to the core node | Implicit: PHS via required sweat rate, with maximum sweat rate rising with metabolic rate (250–400 W·m$^{-2}$); HSDA via empirical sweat-rate equations from required and maximum evaporation |
| ***Respiratory heat loss*** | $\dot{V}_E = \dot{V}O_2$ × ventilatory equivalent (rising above $V_{T1}$, 60% $\dot{V}O_{2max}$) + an additive hyperthermic term in $T_{body}$ (Gibbons et al. 2022a, b), capped at $\dot{V}_{E,max}$ from $\dot{V}O_{2max}$; expired-air temperature from inspired air (Livingstone et al. 1994) | Fanger (1973) form, proportional to metabolic rate: $C_{res} = 0.0014\ M\ (34 - T_a)$, $E_{res} = 0.0173\ M\ (5.87 - p_a)$; losses taken from the airway tissues of the head, neck and thorax (Fiala et al. 1999) | Fanger form on total heat production, $[0.0014\ (34 - T_a) + 0.0173\ (5.87 - p_a)] \times \Sigma Q$, from the chest core; no effect of intensity or body temperature on ventilation | ASHRAE (Fanger) form, proportional to metabolic rate: $C_{res} = 0.0014\ M\ (34 - T_a)$, $E_{res} = 0.0173\ M\ (5.87 - p_a)$ (Vanos et al. 2023, supplementary methods) | Fanger form, proportional to metabolic rate: $0.0014\ M\ (34 - T_a) + 0.0173\ M\ (5.87 - p_a)$ | PHS: $\dot{V}_E = 0.076\ M$ and expired-air temperature from inspired air (Livingstone et al. 1994), so $C_{res}$ and $E_{res}$ scale with $M$ ($E_{res}$ after Varène 1986). HSDA: no respiratory term in required evaporation |
| ***Core temperature sites*** | Central blood node read as $T_{heart}$ (also the control input); passively perfused rectal compartment with exercise-dependent perfusion | Head core (hypothalamic, drives control), thorax core (oesophageal) and abdomen core (rectal) | Central blood node plus segmental artery, vein and core nodes (control from head core); rectal read as pelvis core | Single core temperature; critical limit 43 °C, reached after 17.88 kJ/kg of heat storage | Single core node | PHS: rectal, derived from mean core temperature via an empirical oesophageal–rectal relation. HSDA: single core (rectal) temperature |
| ***Spatial resolution*** | Heart (central blood), lumped core, rectal, active and inactive muscle, fat and skin | 15 elements, multi-layer, with anterior/posterior/inferior sectors | 17 segments, 85 nodes, with arteriovenous anastomoses in hands and feet | Whole-body heat balance (single core compartment with an assumed infinitesimally small skin layer); 3- or 6-h exposures; sun (MRT = Ta + 15 °C) or shade | Two concentric nodes (core, skin shell) | PHS: whole-body heat balance; storage split between core and skin. HSDA: single core, no storage term |
| ***Individual characteristics*** | Age, sex, body composition, $\dot{V}O_2$max, acclimatisation, menstrual phase, burn grafts, spinal cord injury, energy deprivation, load carriage. Can easily accommodate new factor. | One reference person (73.5 kg, 14% fat), unacclimatised; anthropometric individualisation in later work | Height, mass, age, sex, body fat, cardiac index; age modifies sweating and vasodilation (step at 60 y), shivering (decade steps to the 60s) and basal blood flow (steps from 50 y, ×0.7 at ≥ 70 y); cold acclimation via brown adipose tissue | Two female profiles: younger (18–40 y) and older (> 65 y) adults, differing in body size, maximum sweat rate and skin wettedness; unclothed or clothed | Body mass and surface area only | PHS: body size, acclimatisation. HSDA: height, mass, acclimatisation, hydration, initial core and skin temperature |
| ***Physiological outputs available to downstream models*** | Regional blood flows (skin, active and inactive muscle, splanchnic and renal, cerebral) bounded by maximal cardiac output, sweat-derived fluid loss and minute ventilation, allowing direct coupling to cardiovascular and respiratory models | Segmental skin blood flow and sweat rate | Segmental tissue blood flows; not bounded by cardiovascular capacity | Survivability (core 43 °C within 3 or 6 h) and maximum sustainable metabolic rate for liveability. | Whole-body skin blood flow and sweat rate; no cardiac output limit or regional flows | Core (rectal) temperature and sweat rate or water loss only; no blood flows, so no mechanistic link to cardiovascular strain |
| ***Software*** | Julia (ModelingToolkit, OrdinaryDiffEq); proprietary, available for research under licence. | Not openly released | Python 3 (open source) | Python (open source) | Published equations; open Python implementations (e.g. pythermalcomfort) | PHS: full program published in ISO 7933 (Annex E); open implementations (e.g. pythermalcomfort). HSDA: published equations and spreadsheet (Potter et al. 2017) |

ACSM, American College of Sports Medicine; HSDA, Heat Strain Decision Aid; MRT, mean radiant temperature; PAR, physical activity ratio; PHS, Predicted Heat Strain; PMV, predicted mean vote; pa, ambient water vapour pressure; Ta, air temperature; $\dot{V}$E, minute ventilation; $\dot{V}O_2$max, maximal oxygen uptake; VT1, first ventilatory threshold.

**Table S2.** Principal TeMP parameters and their basis.

| Parameter | Value | Basis | Source |
|---|---|---|---|
| **Metabolism** | | | |
| Basal energy expenditure | 0.32 × FFM^0.75 MJ·d⁻¹ | Literature | Pontzer et al. (2021) |
| Age decline in basal energy expenditure | −0.45% per year above 63 y | Literature | Pontzer et al. (2021) |
| Seated resting metabolic rate | 1.3 × basal energy expenditure | Literature | Quiet sitting, 1.3 MET (Ainsworth et al. 2011) |
| Activity metabolic rate | Resting rate + (METs − 1) × 3.5 mL $O_2 \cdot kg^{-1} \cdot min^{-1}$; 20.9 kJ per L $O_2$ | Literature | ACSM metabolic equations (Liguori et al. 2021) |
| Load carriage (walking, running) | Metabolic cost × (body mass + load) / body mass | Derived | ACSM equations; checked against Pandolf et al. (1977) |
| Cycling gross efficiency | 20% | Literature | ACSM leg-cycling equation |
| Exercise metabolic cost and core temperature | × 1.25^(($T_{heart}$ − 37)/10) above 37 °C (≈ 2.3% per °C); exercise metabolism only | Fitted | Exercise validation data; higher $\dot{V}O_2$ as core temperature rises (Fink et al. 1975) |
| Heat acclimatisation, resting metabolic rate | −18% at full acclimatisation | Literature | Lee & Kim (2018) |
| Energy restriction, resting metabolic rate | × 0.90 / 0.75 / 0.60 (mild / moderate / severe) | Assumed | Low resting metabolism and body temperature in anorexia nervosa (Wakeling & Russell (1970)) |
| **Circulation and skin blood flow** | | | |
| Maximal cardiac output (COmax) | 5.5 × $\dot{V}O_{2max}$ ($L \cdot min^{-1}$) + 5 $L \cdot min^{-1}$ | Derived | Linear cardiac output–$\dot{V}O_2$ relation (Åstrand et al. 1964; Hermansen et al. 1970; Beck et al. 2006) |
| Skin blood flow control signal | 0.8 $T_{heart}$ + 0.2 $T_{skin}$ | Assumed | Conventional core weighting (Werner 2010; Wyss et al. 1974) |
| Release of vasoconstrictor tone | Onset 35.86 °C; 0.4 $L \cdot min^{-1} \cdot m^{-2}$ per °C | Fitted | Resting baselines (Che Muhamed et al. 2016; Périard et al. 2012; Meade et al. 2023, 2025) |
| Active vasodilation | Onset 36.51 °C; saturation 38.71 °C | Fitted | Meade et al. (2023, 2025) |
| Exercise, vasodilation onset and saturation | Raised by 0.6 °C × (fraction of $\dot{V}O_2$ reserve − 0.45), above 0.45 | Literature / fitted | Direction from Taylor et al. (1988); magnitude fitted to exercise validation data |
| Skin blood flow ceiling at rest | 24.2% of $CO_{max}$ | Literature | Minson et al. (1998) |
| Skin blood flow ceiling during exercise | Falls to 9% of $CO_{max}$ at $\dot{V}O_{2max}$ (midpoint 60% $\dot{V}O_{2max}$) | Assumed | Redistribution to active muscle (Rowell 1974) |
| Active muscle blood flow, maximum | 60% of $CO_{max}$ | Literature | Mortensen et al. (2005) |
| Visceral (splanchnic and renal) blood flow | Basal ≈ 65% of resting cardiac output (incl. cerebral); maximum thermal reduction 68% | Literature | Rowell (1974); Minson et al. (1998) |
| Age, skin blood flow ceiling | × 0.56 at 70 y | Literature | Minson et al. (1998) |
| **Sweating** | | | |
| Sweating control signal | 0.9 $T_{heart}$ + 0.1 $T_{skin}$ | Literature | Nadel et al. (1971) |
| Sweating onset | 36.75 °C | Fitted | Meade et al. (2023, 2025) |
| Sweating sensitivity | 164 $g \cdot h^{-1} \cdot m^{-2}$ per °C | Fitted | Meade et al. (2023, 2025) |
| Maximum sweat rate | 800 $g \cdot h^{-1} \cdot m^{-2}$ (1.44 $L \cdot h^{-1}$ for 1.8 $m^2$), unacclimatised, $\dot{V}O_{2max}$ 40 | Literature | Havenith (2001) |
| Fitness, maximum sweat rate | × [1 + 0.25 ($\dot{V}O_{2max}$ − 40)/20], $\dot{V}O_{2max}$ limited to 20–60 | Literature | Havenith (2001) |
| Local skin temperature effect on sweating | × exp[($T_{skin}$ − 34)/10] | Literature | Nadel et al. (1971) |
| Sweating efficiency | 1 − $w^2$/2; maximum skin wettedness 0.60 | Literature / fitted | ISO 7933 (2004); Candas et al. (1979); Meade et al. (2025) |
| Hidromeiosis | Sweat decline beginning ≈73 min into full skin wetting (observed 60–80 min) | Literature | Candas et al. (1980, 1983) |
| Exercise, sweating onset | Lowered by 1.5 °C × fraction of $\dot{V}O_2$ reserve | Fitted | Che Muhamed et al. (2016); Périard et al. (2012); Mora-Rodríguez et al. (2010) |
| Exercise, sweating sensitivity | × (1 + 1.5 × fraction of $\dot{V}O_2$ reserve) | Fitted | As above |
| Age, sweating onset | +0.003 °C per year (+0.13 °C at 70 y) | Fitted | Meade et al. (2023); delayed onset with age (Sagawa et al. 1988) |
| **Individual modifiers of thresholds** | | | |
| Heat acclimatisation, sweating onset | −0.4 °C at full acclimatisation | Assumed | Direction from Yamazaki & Hamasaki (2003): −0.24 °C oesophageal after 6 days |
| Heat acclimatisation, sweating sensitivity | 178 $g \cdot h^{-1} \cdot m^{-2}$ per °C (+8%) | Literature | Within the 0–15% reported (Havenith 2001; Roberts et al. 1977; Yamazaki & Hamasaki 2003) |
| Heat acclimatisation, maximum sweat rate | +25% of the reference maximum | Literature | Havenith (2001) |
| Heat acclimatisation, vasodilation onset | −0.25 °C | Derived | Yamazaki & Hamasaki (2003) |
| Luteal phase | Sweating and vasodilation onsets +0.5 °C | Literature | Hessemer & Brück (1985); Stephenson & Kolka (1985) |
| Spinal cord injury | Retained effector function 0.80 / 0.50 / 0.30 (lumbar / thoracic / cervical) | Derived | Absence of afferent $T_{body}$ feedback below lesion |
| Burn injury (grafted skin) | No sweating or active vasodilation over the grafted area | Derived | Absence of sympathetic innervation in grafts |
| **Respiration** | | | |
| Metabolic minute ventilation | $\dot{V}O_2$ × $EqO_2$; $\dot{V}O_2$ from total metabolic rate (20.9 kJ per L $O_2$) | Derived | Ventilation follows $O_2$ uptake rather than metabolic rate per unit surface area |
| Ventilatory equivalent ($EqO_2$) | 25 below $VT_1$, rising linearly to 35 at $\dot{V}O_{2max}$ | Literature / assumed | Sheel & Romer (2012); Norton et al. (1995) |
| First ventilatory threshold ($V_{T1}$) | ~60% $\dot{V}O_{2max}$ | Literature | Sheel & Romer (2012) |
| Maximal minute ventilation | 12.34 + 28 × $\dot{V}O_{2max}$ ($L \cdot min^{-1}$) | Literature | Blackie et al. (1991) |
| Hyperthermic ventilation (additive) | 6.9 $L \cdot min^{-1}$ per °C of $T_{body}$ (0.8 $T_{heart}$ + 0.2 $T_{skin}$) above 37.15 °C, at rest and in exercise | Fitted | Gibbons et al. (2022a, 2022b) |
| Expired air temperature | 28.56 + 0.115 $T_{air}$ + 0.641 $P_{air}$ (°C) | Literature | Livingstone et al. (1994); ISO 7933 (2004) |
| **Heat exchange and body** | | | |
| Body surface area | DuBois form, Kerr constants | Literature | Kerr (1988) |
| Convective heat transfer coefficient | 8.3 v^0.6 $W \cdot m^{-2} \cdot K^{-1}$ | Literature | Mitchell (1974) |

| Parameter | Value | Basis | Source |
|---|---|---|---|
| Clothing insulation and vapour permeability | Permeability index 0.38 | Literature | ISO 9920 (2007); ISO 7933 (2004) |
| Skin thermal conductivity | 0.37 $W \cdot m^{-1} \cdot K^{-1}$ | Literature | McIntosh & Anderson (2010; Buettner 1936 value, 0.376) |
| Muscle thermal conductivity | 0.49 $W \cdot m^{-1} \cdot K^{-1}$ | Literature | McIntosh & Anderson (2010); gives whole-body tissue insulation 0.11 $m^2 \cdot K \cdot W^{-1}$ when vasoconstricted, vs ≈0.10 in vivo (Veicsteinas et al. 1982) |
| Simulation stop (heat stroke) | $T_{rectal}$ 41 °C | Assumed | Arbitrary but generally viewed as an upper limit during passive heating. |

Basis: Literature, value taken from the cited source; Derived, calculated from literature values; Fitted, set against the named validation data; Assumed, modelling assumption. Temperatures for onsets and saturation refer to each effector's control signal. ACSM, American College of Sports Medicine; $CO_{max}$, maximal cardiac output; $EqO_2$, ventilatory equivalent for oxygen; FFM, fat-free mass; MET, metabolic equivalent; $P_{air}$, ambient water vapour pressure (kPa); $T_{air}$, air temperature; $T_{body}$, mean body temperature (ventilatory drive); $T_{heart}$, heart temperature; $T_{rectal}$, rectal temperature; $T_{skin}$, skin temperature; $\dot{V}_E$, minute ventilation; $\dot{V}O_{2max}$, maximal oxygen uptake; $VT_1$, first ventilatory threshold; w, skin wettedness.

**Table S3.** Studies used to calibrate and evaluate TeMP.

| Study | Participants | Protocol | Variables used | Role |
|---|---|---|---|---|
| Che Muhamed et al. (2016) | Young trained men | Treadmill run, 70% $\dot{V}O_{2max}$, 60 min; 31 °C at 23, 52 and 71% RH | $T_{rectal}$, $T_{skin}$, whole-body sweat rate, heat storage | Exercise sweating; exercise heat partition; resting baseline |
| Périard et al. (2012) | Untrained men | Cycling at 60 and 75% $\dot{V}O_{2max}$ in 40 °C, 50% RH; 60% $\dot{V}O_{2max}$ in 18 °C | $T_{rectal}$ and $T_{skin}$, core-to-skin gradient | Exercise sweating; skin blood flow during exercise; resting baseline |
| Mora-Rodríguez et al. (2010) | Trained and untrained men | Cycling at 40, 60 and 80% $\dot{V}O_{2peak}$ in 36 °C, 25% RH, 2.5 m·s⁻¹ airflow | $T_{rectal}$ and $T_{skin}$, sweat rate | Exercise sweating; effect of fitness; resting baseline |
| Meade et al. (2023) | Young (n = 20) and older (n = 39) adults | Seated rest, 9 h, 40 °C, 9% RH | $T_{rectal}$ and $T_{skin}$, evaporative heat loss, heat storage | Sweating and vasodilation onsets; age effect on sweating onset; resting baseline |
| Meade et al. (2025) | Young adults (n = 12) | Seated rest, up to 9 h, 42 °C above and below the critical wet-bulb limit; 28 °C control | $T_{rectal}$ and $T_{skin}$, whole-body sweat rate | Sweating onset; maximum skin wettedness; resting baseline |
| Morris et al. (2019) | Young men (n = 12) | Seated rest, 2 h, 47 °C/14% RH and 40 °C/51% RH, with and without an electric fan | $T_{rectal}$, whole-body sweat rate | Air movement and evaporation |
| Gibbons et al. (2022a) | Young adults (n = 13) | Seated rest, passive heating (water-perfused suit) to +1 and +2 °C $T_{oes}$ | $T_{oes}$, $T_{skin}$, $\dot{V}_E$ | Hyperthermic ventilation (rest) |
| Gibbons et al. (2022b) | Young adults (6 women, 6 men) | Semi-recumbent cycling at 25% maximal aerobic power, with and without heating (water-perfused suit) to +1 and +2 °C $T_{oes}$ | $T_{oes}$, $T_{skin}$, $\dot{V}_E$, $\dot{V}O_2$ | Hyperthermic ventilation (exercise) |
| **Studies setting individual parameters** | | | | |
| Minson et al. (1998) | Young and older men | Passive whole-body heating to the limit of tolerance | Rise in skin blood flow; splanchnic and renal redistribution | Skin blood flow ceiling; age reduction in ceiling; visceral flow reduction |
| McKenna et al. (2023) | Young and older adults | 3 h, 47 °C, 10% RH with activities of daily living | $T_{rectal}$, sweat rate | Age effects on sweating |
| Griggs et al. (2019) | Able bodied, paraplegia and tetraplegia | 37 °C, incremental rises in RH | $T_{gi}$ | Impact of spinal cord injury level on the change in $T_{gi}$ |
| Kolka and Stephenson (1997) | Early follicular vs mid-luteal phase | Exercise at 80% peak aerobic power and ambient temperature of 35°C, 22% RH | $T_{oes}$ | Impact of menstrual phase on $T_{oes}$ during exercise in the heat. |
| Wakeling and Russell (1970) | Anorexic patients before and after treatment | Baseline body temperature | Oral temperature | Baseline body temperature before and after treatment for anorexia nervosa |
| **Independent comparisons (not used for fitting)** | | | | |
| Saltin & Hermansen (1966) | Young adults (n = 7) | Cycling, 26% $\dot{V}O_{2max}$, 60 min, ~20 °C | $T_{rectal}$ | Figure 9 |
| Drinkwater et al. (1976) | Young women (n = 12) | Treadmill walking, ~30% $\dot{V}O_{2max}$, 50 min, 35 °C, 65% RH | $T_{rectal}$ | Figure 9 |
| Meade et al. (2024) | Older adults (mean 72 y) | Seated rest, 8 h, 22, 26, 31 and 36 °C, 45% RH | $T_{rectal}$ | Figure 5 |

$T_{gi}$, gastrointestinal temperature; $T_{oes}$, oesophageal temperature; $T_{rectal}$, rectal temperature; $T_{skin}$, skin temperature; RH, relative humidity; $\dot{V}_E$, minute ventilation; $\dot{V}O_{2peak}$, peak oxygen uptake; $\dot{V}O_{2max}$, maximal oxygen uptake. Calibration studies were run with each paper's protocol and scored against the reported variability.
